# Fast estimation of genetic correlation for Biobank-scale data

**DOI:** 10.1101/525055

**Authors:** Yue Wu, Kathryn S. Burch, Andrea Ganna, Päivi Pajukanta, Bogdan Pasaniuc, Sriram Sankararaman

## Abstract

Genetic correlation is an important parameter in efforts to understand the relationships among complex traits. Current methods that analyze individual genotype data for estimating genetic correlation are challenging to scale to large datasets. Methods that analyze summary data, while being computationally efficient, tend to yield estimates of genetic correlation with reduced precision. We propose, SCORE, a randomized method of moments estimator of genetic correlation that is both scalable and accurate. SCORE obtains more precise estimates of genetic correlations relative to summary-statistic methods that can be applied at scale achieving a 50% reduction in standard error relative to LD-score regression (LDSC) and a 26% reduction relative to high-definition likelihood (HDL) (averaged over all simulations). The efficiency of SCORE enables computation of genetic correlations on the UK biobank dataset consisting of ≈ 300K individuals and ≈ 500K SNPs in a few hours (orders of magnitude faster than methods that analyze individual data such as GCTA). Across 780 pairs of traits in 291, 273 unrelated white British individuals in the UK Biobank, SCORE identifies significant genetic correlation between 200 additional pairs of traits over LDSC (beyond the 245 pairs identified by both).

## Introduction

Genetic correlation is an important parameter that quantifies the genetic basis that is shared across two traits. Estimates of genetic correlation can reveal pleiotropy, uncover novel biological pathways underlying diseases, and improve the accuracy of genetic prediction [1].

While traditionally reliant on family studies, the availability of genome-wide genetic data has led to several approaches to estimate genetic correlation from these datasets [1]. An important class of methods for estimating genetic correlation relies on computing the restricted maximum likelihood within a bi-variate linear mixed model (LMM), termed genomic restricted maximum likelihood (GREML)[2–5]. However, current GREML methods are computationally expensive to be applied to large-scale datasets such as the UK Biobank [6].

While GREML methods need individual-level data, several methods [7–12], such as LD-score regression (LDSC) [7], have been proposed for estimating genetic correlation using GWAS summary statistics. While methods such as LDSC often have substantially reduced computational requirements relative to GREML, LDSC estimates tend to have large standard errors which increase further when there is a mismatch between the samples used to estimate summary statistics and the reference datasets that are used to estimate LD scores [13]. High-definition likelihood (HDL) [12], a more recent summary-statistic based method, has been shown to be more precise relative to LDSC. HDL, however, requires computing a singular-value decomposition (SVD) of the LD matrix which increases its runtime. Further, recent studies [14, 15] have shown that the accuracy of genetic correlation estimates can deteriorate when there is a mismatch between reference and sample data. Thus, it is critical to develop methods for estimating genetic correlation that can work directly with large individual-level datasets.

We propose, SCORE (SCalable genetic CORrelation Estimator), a randomized Method-of-Moments (MoM) estimator of genetic correlations among traits using individual genotypes that can scale to the dataset sizes typical of the UK Biobank. While SCORE can estimate the heritability of traits as well as the genetic correlation between pairs of traits, we focus on the problem of estimating genetic correlation in this work. SCORE avoids the explicit computation of the genetic relationship matrix (GRM). Instead, we show that the genetic correlation can be computed using a *sketch* of the genotype matrix, *i.e.,* by multiplying the genotype matrix with a small number of random vectors.

In simulations, we show that SCORE yields accurate estimates of genetic correlation across a range of genetic architectures and sample overlap settings. Relative to summary-statistic methods that can be applied to Biobank-scale data, SCORE obtains a reduction in the standard error of 50% relative to LDSC and 26% relative to HDL (averaged across all simulations). Further, SCORE can estimate genetic correlation on ≈ 500K SNPs in ≈ 300K unrelated white British individuals in a few hours, orders of magnitude faster than methods that rely on individual data (GCTA-GREML and GCTA-HE). Analyzing 780 pairs of traits in 291,273 unrelated white British individuals in the UK Biobank, the estimates of genetic correlation at 459, 792 common SNPs obtained by SCORE are largely concordant with those from LDSC (Pearson correlation *r* = 0.95). While 245 pairs of traits are identified to have significant genetic correlation by both methods (using a Bonferroni correction for the number of pairs of traits tested), the reduced standard error of estimates from SCORE leads to the discovery of the significant genetic correlations between additional 200 pairs of traits relative to LDSC. Finally, SCORE detects a significant positive correlation between serum liver enzyme levels (alanine (ALT) and aspartate aminotransferase (AST)) and coronary artery-disease related traits (angina and heart attack) suggesting that coronary artery disease and liver dysfunction harbor a shared genetic component.

## Results

### Methods overview

We assume a pair of traits, ***y*_1_**, ***y*_2_** related to standardized genotypes ***X*_1_, *X*_2_** by the following model:

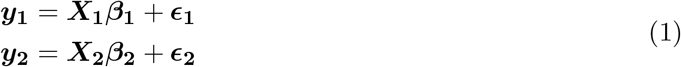

Here we have *N*_1_ samples for trait 1 and *N*_2_ samples for trait 2 of which *N* samples (*N* ≤ *N*_1_, *N* ≤ *N*_2_) contain measurements for both the traits. ***X*_1_, *X*_2_** are the *N*_1_ × *M* and *N*_2_ × *M* matrices of standardized genotypes. ***β*_1_, *β*_2_** are the vectors of SNP effect sizes while ***ϵ*_1_** and ***ϵ*_2_** denote trait-specific environmental noise that is independent of the genetic effect.

Additionally, we assume the effect sizes for the two traits have a mean of 0 and covariance:

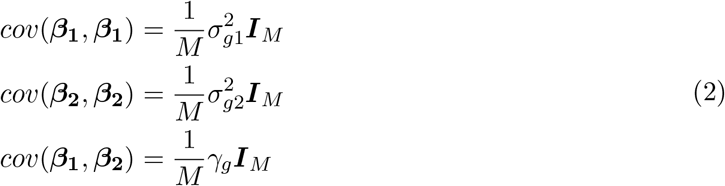

Here ***I***_*M*_ is an *M × M* identity matrix and 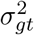 denotes the genetic variance associated with trait *t* (*t* ∈ {1,2}), and *γ_g_* denotes the genetic covariance. The trait-specific environmental noise in each individual is assumed to have zero mean and variance 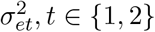 for trait *t*. Further, we assume that traits that are measured on the same individuals have additional environmental covariance *γ_e_*. The genetic correlation parameter *ρ_g_* is defined as 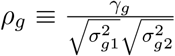. Importantly, SCORE does not make specific assumptions on the distribution of the genetic effect sizes or the environmental noise.

SCORE uses a scalable method-of moments estimator for the genetic correlation, 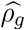 (While SCORE can jointly estimate the genetic and environmental variance and covariance parameters, we focus on the estimates of genetic correlation in this work). SCORE works by finding values of the model parameters i.e., the genetic and environmental variance and covariances, such that the population and sample moments match. The scalability of SCORE arises from the use of a randomized algorithm that operates by multiplying the input genotype matrix with a small number of random vectors thereby avoiding explicitly computing the genetic relatedness matrix (GRM) for each trait. The model described here considers the general case where the two traits are not measured on the same set of individuals (so that it can handle partial or no overlap). When two traits are measured on the same set of individuals (100% sample overlap), further computational efficiency can be obtained for estimating the genetic correlation. We refer to this variant of SCORE as SCORE-OVERLAP (see Supplementary Information). Further, SCORE uses a streaming algorithm that has scalable memory requirements and uses an efficient block Jackknife with a block size of 4000 SNPs to estimate standard errors with little additional computational overhead.

### Accuracy and robustness of SCORE in Simulations

We performed simulations to compare the accuracy of SCORE to other estimators of genetic correlation under different genetic architectures. Specifically, we compared SCORE to methods that use individual data (bi-variate GREML [2], bi-variate Haseman-Elston regression) and methods that rely on summary statistics (LD-score regression (LDSC) [7] and HDL [12]). Bi-variate GREML (GCTA-GREML) and Haseman-Elston regression (GCTA-HE) are implemented in the GCTA software. LDSC is a widely used method to estimate genetic correlation when only summary statistics from GWAS on pairs of traits are available. HDL is a recent summary-statistics based method that has been shown to obtain improved statistical efficiency relative to LDSC.

We performed simulations to assess the estimation accuracy of each method using a subset of 5,000 unrelated white British individuals in the UK Biobank so that all the methods could be run in a reasonable time. Our simulations used 305,630 SNPs with minor allele frequency (MAF) above 1% (we chose these SNPs since these were also used for benchmarking the HDL [12] method). Given the genotypes, we simulated pairs of traits with known values of heritability and genetic correlation under both infinitesimal and non-infinitesimal architectures. Specifically, we varied the heritability of the pairs of traits which we refer to as low-low, low-mid, mid-mid, and high-high heritability settings).

The simulations assume that the two traits are measured on the same set of individuals so that both SCORE and SCORE-OVERLAP can be applied in this setting. We confirmed that SCORE (with *B* = 10 and *B* = 100 random vectors) and SCORE-OVERLAP yield nearly identically results across the 32 architectures (Supplementary Table S1). In all our results, we use SCORE with *B* = 10 as our default. Across these architectures, the standard error (SE) of SCORE range from 0.89 to 1.17 relative to SE of GCTA-GREML with the SE of SCORE being 2.5% higher than that of GCTA-GREML on average Figures 1, 2). Interestingly, GCTA-HE tends to have a SE of 1.38 times that of SCORE on average (range 1.2 to 1.6). Compared to methods that rely on summary statistics, the SE of LDSC relative to SCORE is 2 on average (range 1.26 to 2.63) while the SE of HDL relative to SCORE is 1.36 (range 1.05 to 1.65) (Supplementary Table S2). The reduction in the SE of SCORE relative to the summary statistic-based methods is equivalent to a three-fold increase in sample size over LDSC and a 85% increase in sample size over HDL on average (the bias, SE and MSE of each of the methods is listed in Supplementary Tables S3, S4).

**Figure 1:**
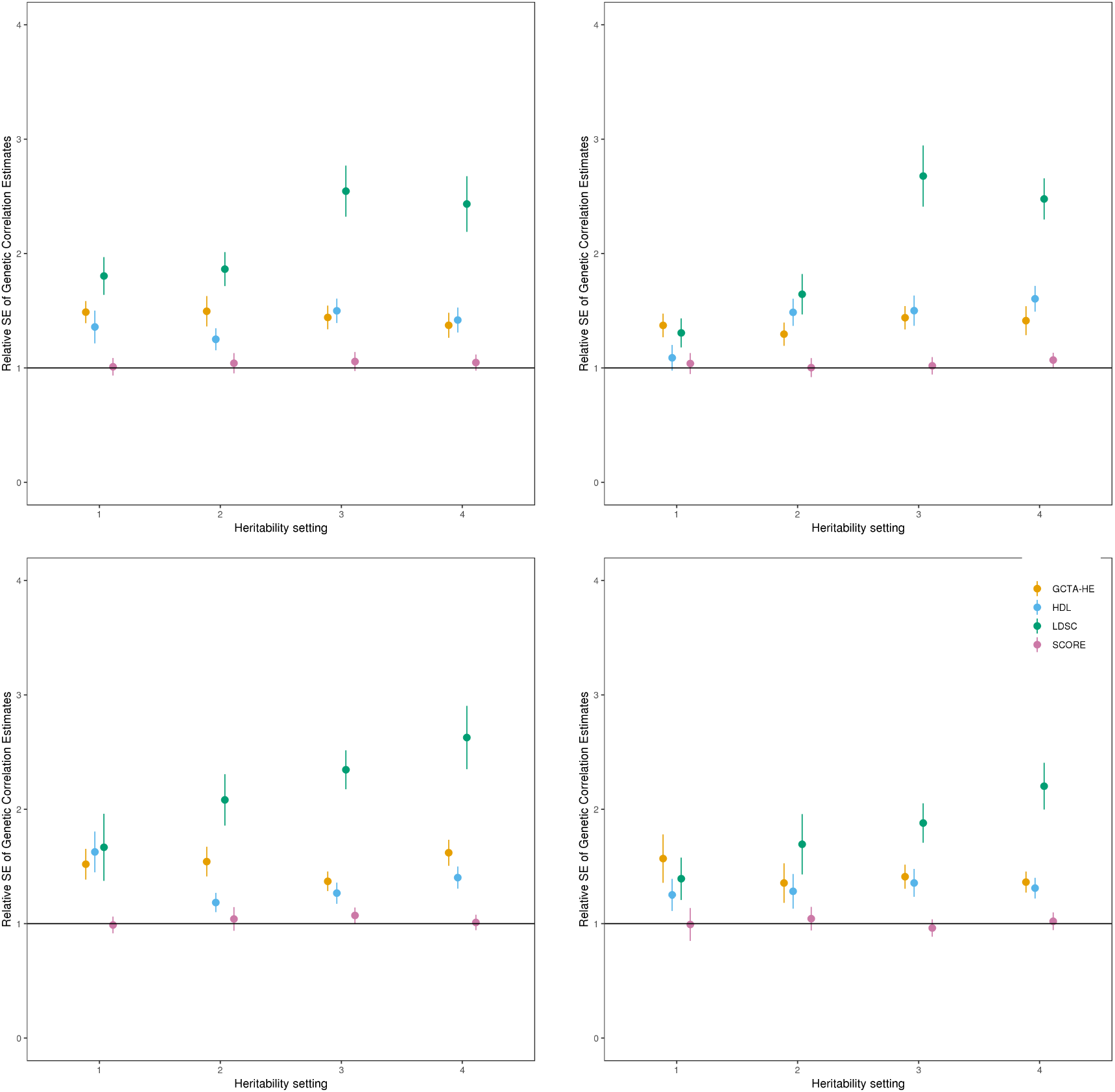
Comparison of the estimates of genetic correlation from SCORE with GCTA-GREML, GCTA-HE, LDSC, and HDL in small-scale simulations (*N* = 5,000 unrelated individuals, *M* = 305,630 SNPs) We simulated pairs of phenotypes under 16 different infinitesimal genetic architectures. Each panel corresponds to a different value of the genetic correlation chosen from the set: {0, 0.2, 0.5, 0.8}. Within each panel, we varied the SNP heritability for the pair of traits across {(0.1, 0.2), (0.2, 0.6), (0.5, 0.5), (0.6, 0.8)}. We plot the standard error (SE) of each method relative to GCTA-GREML. We ran LDSC with in-sample LD and HDL with eigenvectors that preserve 90% variance. We estimate the standard error of the relative SE using Jackknife (error bars denote 1 standard error).

**Figure 2:**
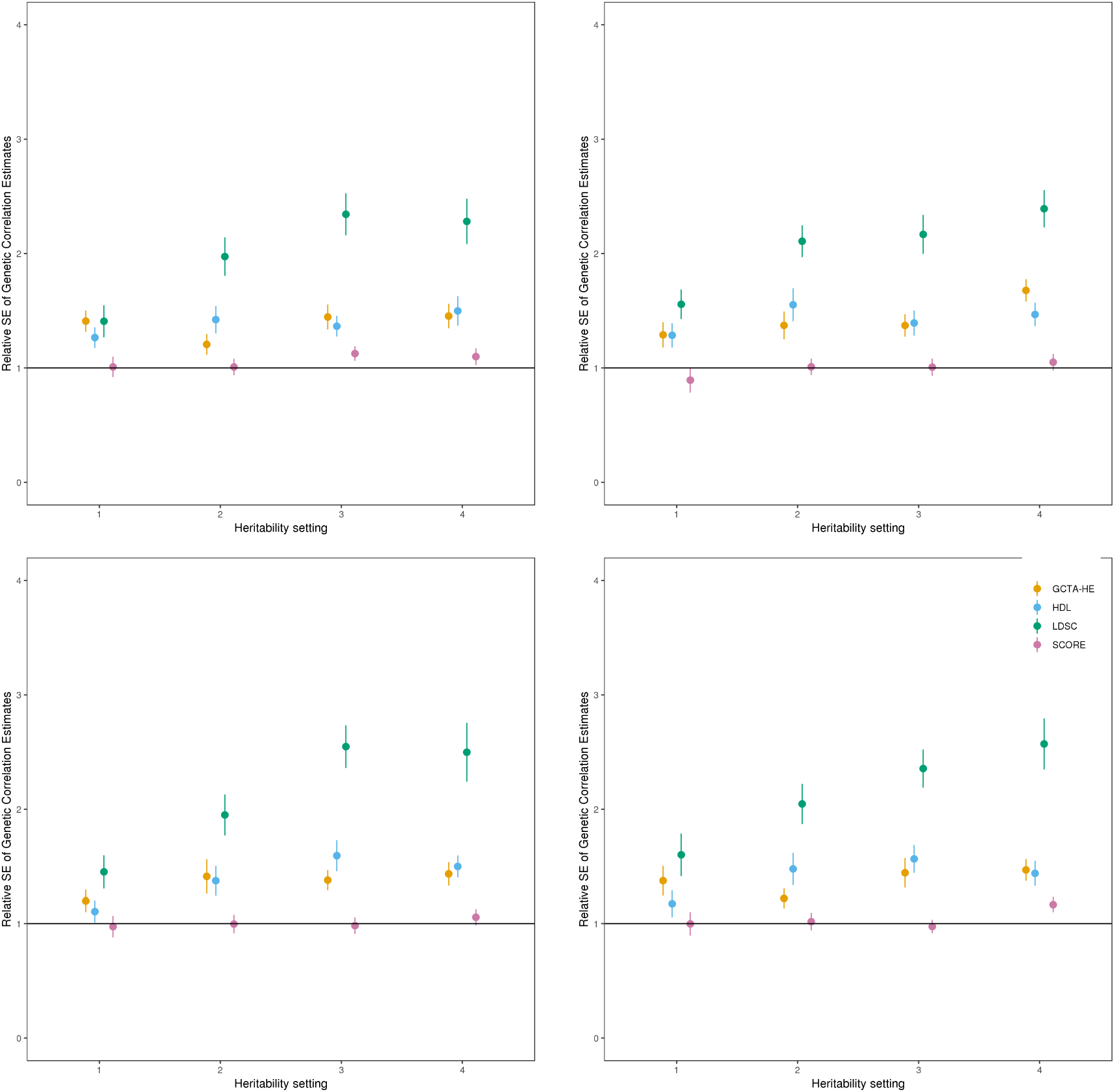
Comparison of the estimates of genetic correlation from SCORE with GCTA-GREML, GCTA-HE, LDSC, and HDL in small-scale simulations (*N* = 5,000 unrelated individuals, *M* = 305,630 SNPs). We simulated pairs of phenotypes under 16 different non-infinitesimal genetic architectures. Each panel corresponds to a different value of the genetic correlation at SNPs causal for both traits: {0, 0.2, 0.5, 0.8}. The causal variants are distributed uniformly across the genome. Within each panel, we varied the per-SNP heritability of variants causal for both traits to be proportional to {(0.1, 0.2), (0.2, 0.6), (0.5, 0.5), (0.6, 0.8)}. We plot the SE of each method relative to GCTA-GREML. We ran LDSC with in-sample LD and HDL with eigenvectors that preserve 90% variance. We estimate the standard error of the relative SE using Jackknife (error bars denote 1 standard error).

We performed additional simulations to investigate the robustness of SCORE. We investigated the impact of sample overlap under an infinitesimal genetic architecture with true genetic correlation *ρ_g_* = 0.5. We observe that the MSE of SCORE to GCTA-GREML and LDSC remains stable as a function of sample overlap (Supplementary Figure S1 and Supplementary Table S5 for the bias, SE, and MSE of SCORE, GCTA-GREML, and LDSC as a function of sample overlap). We also verified that the Jackknife standard error estimate used in SCORE is generally accurate while being conservative for low heritabilities (Supplementary Table S6). Finally, we evaluated the accuracy of SCORE when applied to pairs of traits where one of the traits is binary while the other is continuous. We simulated continuous traits and binary traits where we varied the prevalence of cases in the binary trait. We observe that the standard error of the estimates of genetic correlation obtained by SCORE tend to have relatively low standard error provided the prevalence of the trait is greater than 0.5% (Supplementary Table S7). Thus, we recommend applying SCORE to traits whose prevalence is no less than 0.5%

### Computational Efficiency

We investigated the computational efficiency of SCORE relative to GCTA-GREML and GCTA-HE. The runtime and memory usage of summary statistic methods (LDSC and HDL) depends on the time needed to compute LD scores and summary statistics of each trait. In addition, HDL also requires the computation of the singular value decomposition (SVD) of LD matrices which is a computationally expensive step. Thus, we do not include runtimes for LDSC and HDL in these comparisons. We varied the number of individuals while the number of SNPs was fixed at 459, 792. Figure 3 shows that GCTA-GREML and GCTA-HE could not scale beyond sample sizes greater than 100,000 due to the requirement of computing and operating on a GRM (we extrapolate the runtime of GCTA-GREML and GCTA-HE to be about 340 days and 44 days on the set of 291,273 unrelated white British individuals in the UK Biobank). On the other hand, SCORE ran in about 1.5 hours on the set of 291, 273 individuals using partial overlap mode with *B* = 10 random vectors while the SCORE-OVERLAP variant ran in about 1 hour on the same dataset.

**Figure 3:**
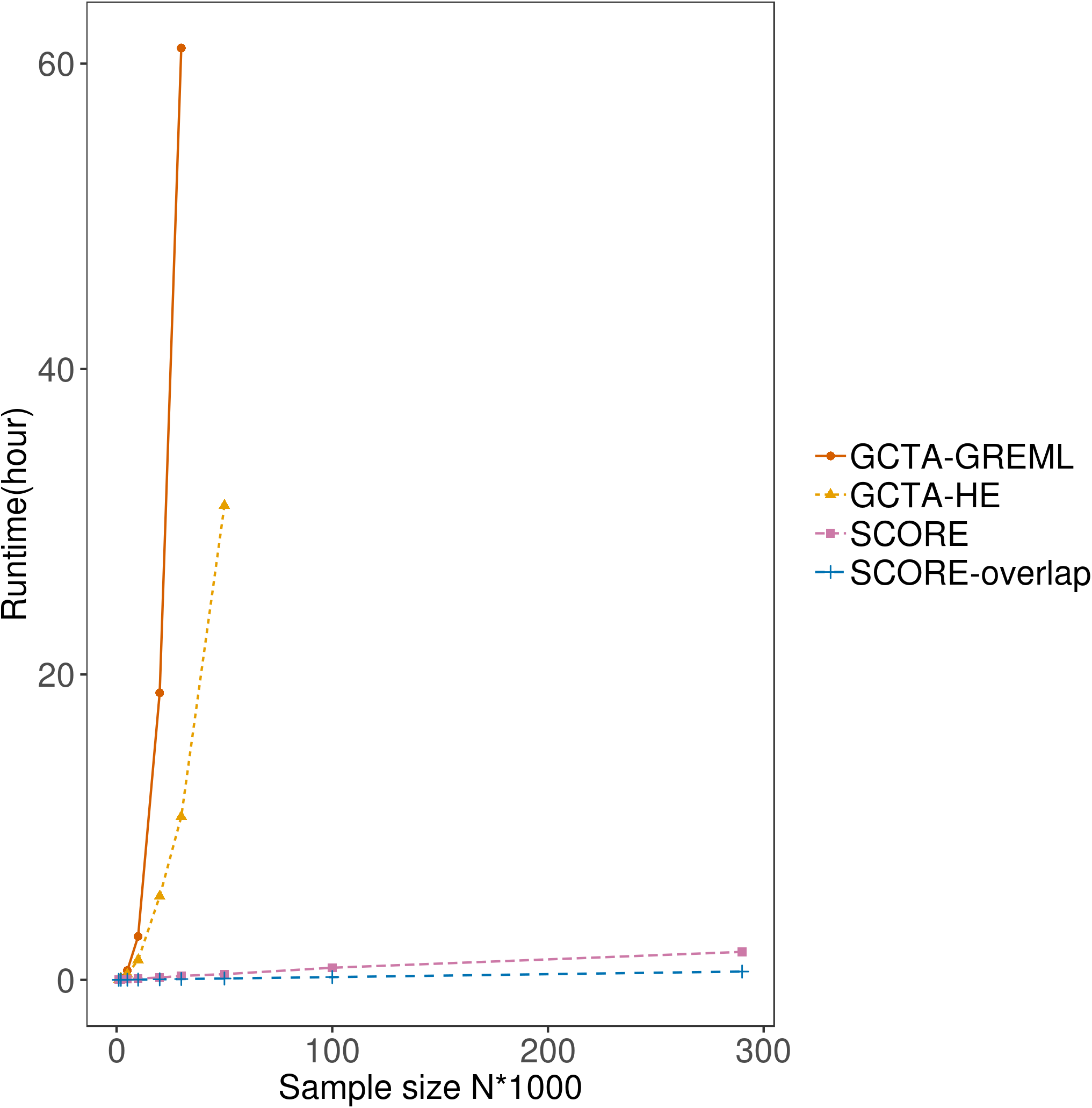
Comparison of the runtime of SCORE with GCTA-GREML and GCTA-HE as a function of the number of samples. The samples were obtained as subsets of unrelated, white British individuals in the UK Biobank. We plot the runtime of both SCORE (that can handle any degree of sample overlap) and its variant, SCORE-OVERLAP, that is designed for 100% sample overlap. SCORE runs in a few hours on the largest dataset of 276, 451 individuals and 459, 792 SNPs.

### Application of SCORE to UK Biobank

We applied SCORE to estimate genetic correlations across phenotypes in the UK Biobank. We restricted our analysis to SNPs on the UK Biobank Axiom array, filtering out markers that were not in HWE, had high missingness rate (> 1%), or low minor allele frequency (< 1%). We also restricted our analysis to unrelated individuals with self-reported white British ancestry. After quality control, we obtained 276,451 individuals and 459,792 SNPs (Methods). We chose traits that have missingness < 30% and disease traits with prevalence larger than 0.5% resulting in a total of 40 phenotypes consisting of 14 binary traits, 3 categorical traits, and 23 continuous traits. The 40 phenotypes could be classified into nine groups: glucose metabolism and diabetes, socioeconomic and general medical information, environmental factor, coronary artery disease related, autoimmune disorders, psychiatric disorders, anthropometric, blood pressure and circulatory, and lipid metabolism (Supplementary Table S8).

We compared the estimates of genetic correlation obtained by LDSC versus SCORE for a subset of 28 traits, in which LDSC produced valid estimates of genetic correlation, *i.e*., traits for which none of the genetic correlation estimates were NA (Figure 4). While the point estimates of genetic correlation from the two methods are highly concordant (Pearson correlation r = 0.95), the SE of LDSC is about 1.57 times that of SCORE on average which is equivalent to a 2.46-fold increase in sample size using SCORE (see Supplementary Figures S2,S3). In total, 192 pairs of traits were detected to have a significant non-zero genetic correlation by both SCORE and LDSC after Bonferroni correction for all pairs across the original set of forty phenotypes 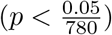. Consistent with its reduced SE, SCORE found 58 pairs with significant genetic correlation after Bonferroni correction that were not detected as significant by LDSC (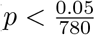; stars in Figure 4). We conclude that SCORE obtains improved power to identify statistically significant genetic correlations relative to LDSC.

**Figure 4:**
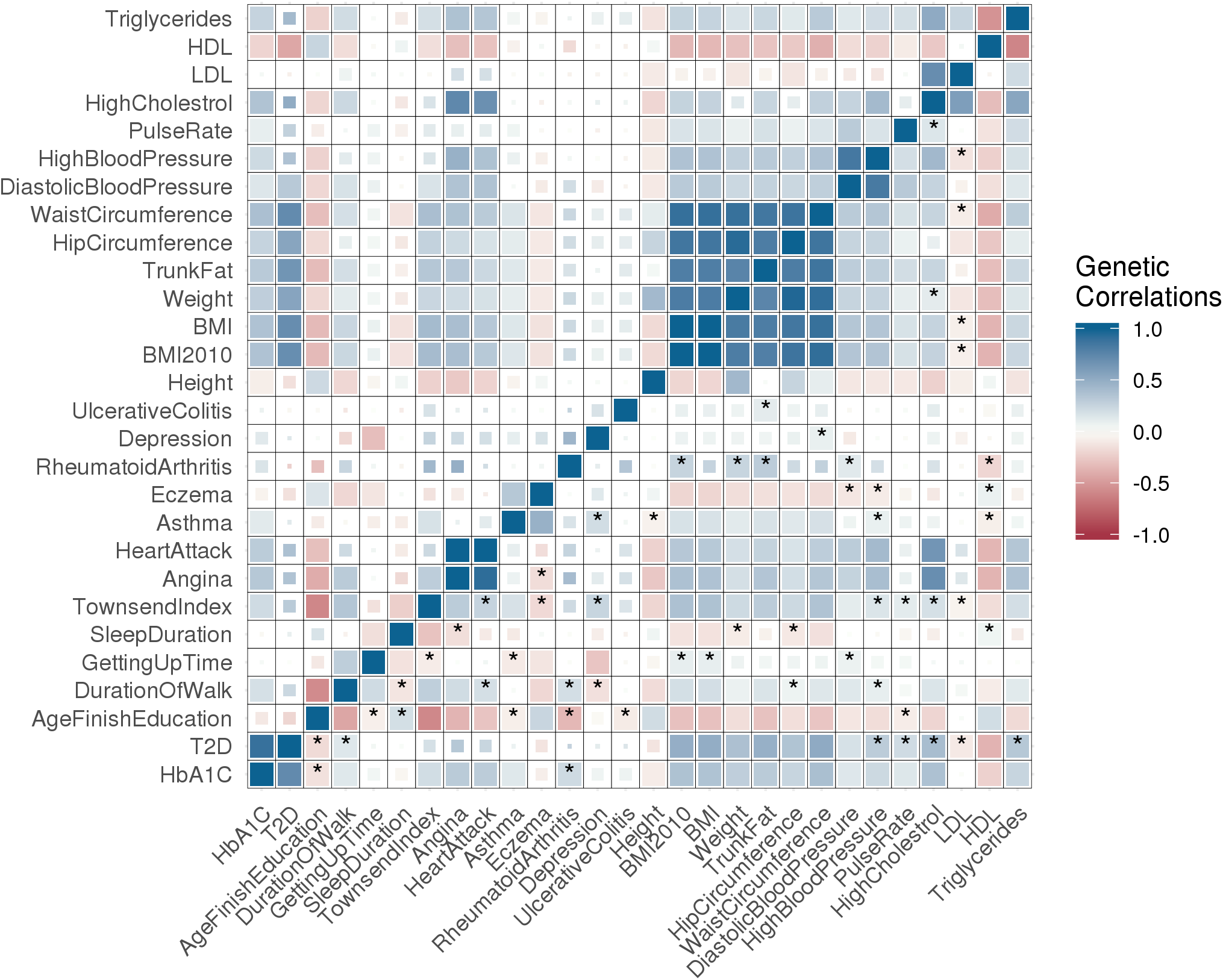
Genetic correlation estimates in the UK Biobank: We plot the genetic correlation estimates from SCORE (bottom triangle) and LDSC (upper triangle) across pairs of 28 phenotypes. Larger filled squares correspond to significant pairs after Bonferroni correction at a 5% significance level while smaller squares correspond to pairs that are significant at a 5% significance level but are not significant after accounting for multiple testing. Star indicates pairs that are found to be significant by SCORE but not by LDSC.

We obtain concordant results when analyzing all pairs in our initial set of forty traits. While the point estimates of SCORE and LDSC are highly correlated (Pearson correlation *r* = 0.96), the SE of LDSC is about 1.8 times that of SCORE on average, equivalent to a 3.24-fold increase in the sample size. In this setting, SCORE found 200 additional pairs of traits over LDSC (beyond the 245 pairs identified by both) while LDSC detected one pair as significant that SCORE did not detect as significant. The estimates of SCORE for all 40 traits are shown in Supplementary Figure S4.

To gain further insights into SCORE, we examined the SE of genetic correlation estimates for pairs of traits according to whether the traits were both binary, both quantitative, or had one member of the pair being binary while the other was quantitative. The SE is largest when both traits are binary, intermediate when one of the traits is binary, and lowest when both traits are quantitative (average SE 0.082, 0.035, and 0.02 respectively; Supplementary Figure S5). We note that the SE increases when the prevalence of the binary trait decreases: the mean SE is 0.017 when the binary trait has prevalence > 0.25 while the mean SE is 0.047 for pairs in which the binary trait has prevalence < 0.05 (Supplementary Figure S6).

To further illustrate its utility, we applied SCORE to estimate genetic correlation between coronary artery-disease related traits included in our set of forty traits (angina and heart attack) and serum biomarkers (alanine (ALT) and aspartate aminotransferase (AST)). It has been shown that serum liver enzyme levels, including ALT and AST, are markers of liver health and hepatic dysfunction and are associated with coronary heart diseases [16–18], though the strength and consistency has varied among the studies [16]. We observed significant positive genetic correlations between ALT/AST and the two coronary artery-disease related trait (0.257± 0.04 and 0.169 ±0.032 for angina with ALT and AST respectively; 0.239 ± 0.053 and 0.148 ± 0.04 for heart attack with ALT and AST respectively). Our novel finding of significant positive genetic correlations suggests hepatic dysfunction (higher serum levels of ALT and AST) and coronary artery disease have a shared genetic component [16–18].

## Discussion

We have described SCORE, a scalable and accurate estimator of genetic correlation. We observe that the estimates of genetic correlation obtained by SCORE obtain accuracy comparable to GREML [13] while being scalable to Biobank-scale data. SCORE can estimate the genetic correlation across pairs of traits when applied to ≈ 500K common SNPs measured on ≈ 300K unrelated white British individuals in the UK Biobank within a few hours. Compared to summary-statistic methods, SCORE obtains a reduction in the average standard error of 61% relative to LDSC and 24% relative to HDL. In application to 780 pairs of traits in the UK Biobank, SCORE discovered 200 pairs of traits with significant genetic correlation (after correcting for multiple testing) that were not discovered by LDSC.

We discuss several limitations of SCORE. Firstly, SCORE requires access to individual genotype and trait data. Summary-statistic methods such as LDSC and HDL have the advantage of being applicable in settings where access to individual-level data can be challenging. While summary-statistic methods also have the advantage of being relatively efficient, it is important to keep in mind that the summary statistics are dependent on specific choices of marker sets and covariates. Applying these methods to different sets of covariates and marker sets requires regenerating the summary statistics (and auxiliary information such as LD score matrices). Second, the model underlying SCORE assumes a quantitative trait. We have shown empirically that SCORE provides accurate estimates of genetic correlation when applied to binary traits provided the traits are not too rare (prevalence > 0.5%). It would be of interest to extend SCORE to the setting of binary traits along the lines of the PCGC method [11]. Finally, while SCORE estimates genome-wide genetic correlation, efficient methods that can partition genetic correlation across genomic annotations can provide novel insights into the shared genetic basis of traits.

## Methods

We describe our model in the general setting, where the traits are not observed on the same set of individuals. Assume we have *N*_1_ individuals for trait 1 and *N*_2_ individuals for trait 2 of which *N* individuals (*N* ≤ *N*_1_,*N* ≤ *N*_2_) contain measurements for both the traits. We have defined ***X*_1_, *X*_2_** to be the *N*_1_ × *M* and *N*_2_ × *M* matrices of standardized genotypes obtained by centering and scaling each column of the unstandardized genotype: ***G_1_*** and ***G_2_*** so that ∑_*n*_ *x_t,n,m_* = 0 for all *m* ∈ {1,…,*M* },*t* ∈ {1,2}. Let ***y*_1_, *y*_2_** denote the two vectors of phenotypes with size *N*_1_ and *N*_2_ respectively. Additionally, we define an *N*_1_ × *N*_2_ indicator matrix, ***C*** where ***C***_*ij*_ = 1 when individual *i* among samples measured for the first phenotype and *j* in samples measured for the second phenotype refer to the same individual and 0 otherwise. We define ***β*_1_, *β*_2_** to be vectors of SNP effect sizes of length *M*.

We assume the following model relating a pair of traits ***y*_1_, *y*_2_**:

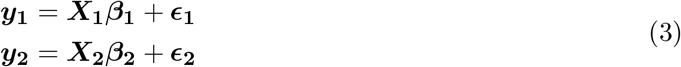

We assume 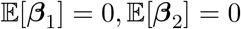, and have the covariance matrix:

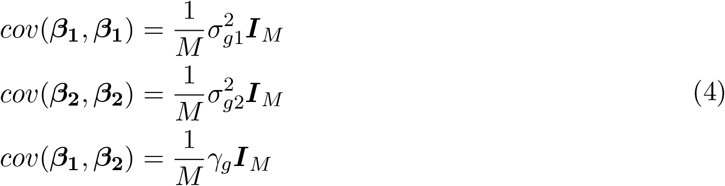

For the environmental effects, we assume 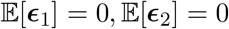, and have the covariance matrix:

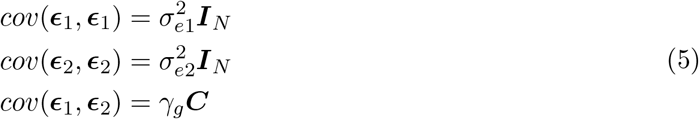

SCORE does not make additional assumptions on the distribution of the genetic effect sizes or the environmental noise.

### Method of Moments (MoM)

SCORE uses a Method of Moments (MoM) estimator to estimate the parameters: 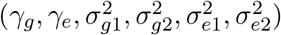. Since the mean of ***y*_1_** and ***y*_2_** are zero, we focus on the covariance. The population covariance of the concatenated phenotypes 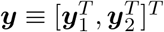 is now:

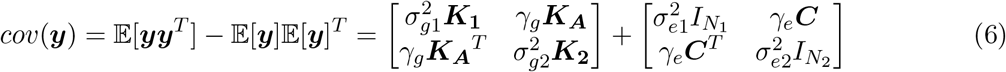

Here 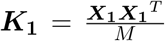 is the GRM for the samples observed for the first trait while 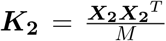 is the GRM for the samples for the second trait and ***K_A_*** is the GRM for pairs of samples across traits: 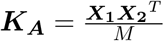.

The MoM estimator is obtained by equating the population covariance to the empirical covariance, estimated by ***yy^T^***:

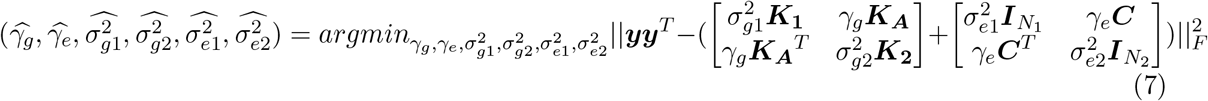

Let 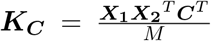. The MoM estimator for genetic covariance satisfies the set of normal equations:

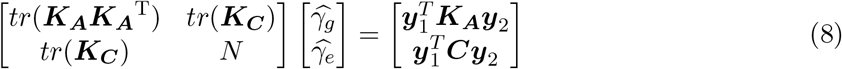

Given each of the coefficients of the normal equations, we can solve analytically for 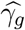, and 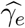.

The variance components 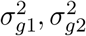 can be estimated with the same idea as described previously [19, 20]. The MoM estimate of the genetic correlation is given by the plug-in estimate:

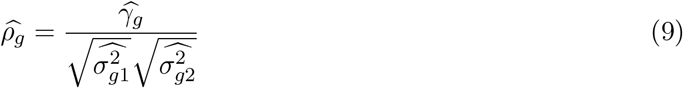

### SCORE: Scalable genetic Correlation Estimator

Naive computation of the MoM estimate requires computing tr(***K_A_K_A_***^T^) which requires 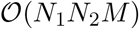 operations, where *N*_1_, *N*_2_ are the sample size of each of the traits. To overcome this computational bottleneck, we replace *tr*(***K_A_K_A_***^T^) with an unbiased randomized estimate: 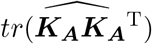 [21].

Given B random vectors, 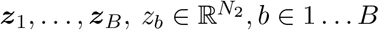 drawn independently from a distribution with zero mean and identity covariance matrix ***I***_***N_2_***_, our estimator is given by:

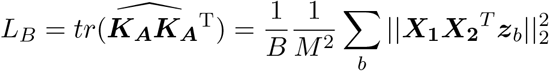

The SCORE estimator 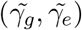 is obtained by solving Equation 8 by replacing *tr*[***K_A_K_A_***^T^| with *L_B_A__*.

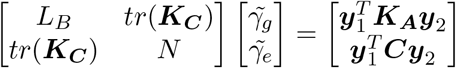

Here *tr*(***K***_*C*_) denotes the sum of the squared genotypes for individuals measured on both traits so that the computation complexity of *tr*(***K***_*C*_) is 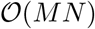. Computing *L_B_* requires multiplying the genotype matrices **X**_1_ and **X**_2_ with *B* vectors resulting in a runtime of 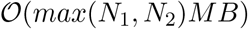. Leveraging the fact that each element of the genotype matrix takes values in the set {0,1,2}, *L_B_* can be computed in time 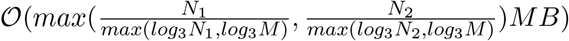 [22].

## Simulations

### Simulations to assess accuracy

We performed simulations on a subset of 5, 000 unrelated white British individuals from the UK Biobank so that all methods compared could be run in a reasonable time. Our simulations used 305, 630 SNPs with minor allele frequency (MAF) above 1% (we chose these SNPs since these were also used for benchmarking the HDL [12] method).

Given the genotypes, we simulated a pair of traits assuming infinitesimal genetic architectures. We varied the genetic correlation across {0, 0.2, 0.5, 0.8} and the heritability of the pair of traits across values of {(0.1, 0.2), (0.2, 0.6), (0.5, 0.5), (0.6, 0.8)} corresponding to the situation where both traits have low heritability, one trait has low heritability while the other has moderate heritability, both traits have moderate heritability and both have high heritability. We assume complete sample overlap and no environmental correlation, and the environmental variance is set so that the trait variance is 1. We simulated a total of 100 replicates for each combination of trait heritabilities and genetic correlation.

In our next set of simulations, we simulated pairs of traits assuming non-infinitesimal genetic architectures. For each genetic variant *m*, we specify a causal status, *c_m_*, which is a 2 × 1 indicator vector with entries taking values in {0,1}. We fix the probability 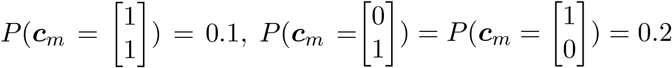, and a genetic variant has no effect on any trait with probability of 0.5. The effect size ***β***_*m*_ for genetic variant *m* on both traits are drawn from the following distribution:

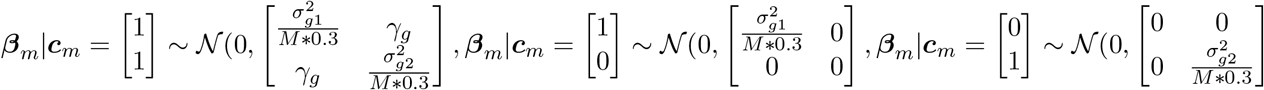

We vary *γ_g_* over the set of values: {0,0.2,0.5,0.8}. Under this model, the true total expected genome wide genetic correlation is {0, 0.06, 0.15, 0.24}. We assume complete sample overlap and no environmental correlation. We simulated a total of 100 replicates for each combination of trait heritabilities and genetic correlation.

### Data processing

For simulations, the LD scores are based on the 305, 630 SNPs chosen for the simulations. The LD scores are generated from a random subset of 50,000 individuals in the UK Biobank (the 5,000 individuals used in our simulations were a subset of the 50, 000 individuals used to compute LD score). For analysis of UK Biobank traits, the LD scores are based on 459, 792 SNPs. The LD scores were computed using flags *l*2 and *ld* -- *wind* -- *kb*2000.0.

The UK Biobank summary statistics input to LDSC were generated using PLINK. We used linear regression to generate summary statistics for continuous traits and categorical traits and logistic regression for binary traits. In computing summary statistics, we include the following covariates: age, gender, principal components 1-10, assessment center, and genotype measurement batch. We used the same covariates as input to SCORE.

We ran LDSC under default settings with an unconstrained intercept. For HDL, we used the eigenvectors of the LD matrix released by HDL which preserve 90% variance of the LD blocks as recommended by the study authors. This computation used the same set of genetic variants as our simulations and 336, 000 samples [12].

### Simulations to assess the impact of sample overlap

We fixed the true heritability for the pair of traits to {0.2,0.6}, and the true genetic correlation to 0.5. We assume there is no environmental correlation. Given the genotype, we simulated a pair of traits under an infinitesimal model. For each trait, we fixed the sample size to 5000 and varied the proportion of sample overlap across the values {0, 0.2, 0.5, 0.8, 1} (ranging from no overlap to complete overlap). Specifically, for overlap proportion equal to 0, we have 5000 samples with observations on the first trait and a distinct set of 5000 samples with observations on the second trait. Similarly, for overlap proportion equal to 1, we have 5000 samples with observations on both traits. For each value of sample overlap, we simulated 100 replicates. We estimated genetic correlation with SCORE, LDSC, and GCTA-GREML. We plot the relative SE in Figure S1 and report the bias, MSE and variance in Table S5.

### Simulations to assess the impact of binary traits

We performed simulations to assess the accuracy of SCORE on binary traits. Given the unrelated white British 276,451 individuals in the UK Biobank measured on 459,792 genetic variants, we fixed the true heritability of the pair of traits to {0.272, 0.12} and the true genetic correlations to —0.23. We also varied the environmental correlation across the values in the set {0.04, —0.04, 0}. For each configuration, we simulated 100 replicates and estimated genetic correlation with SCORE.

To simulated binary traits, we picked the second trait and converted it to a binary trait by thresholding the underlying continuous trait such that the prevalence varied across {0.01%, 0.5%, 1%}. We report the mean and the standard error of the estimates of SCORE across 100 replicates as a function of the prevalence.

### Quality control of UK Biobank genotype array data

We restricted our analysis to SNPs genotyped on the UK Biobank Axiom array, filtering out markers that had high missingness rate (> 1%) and low minor allele frequency (< 1%). Moreover, SNPs that fail the Hardy-Weigerg test at significance threshold 10^-7^ were removed. We also filter the samples that have a genetic kinship with any other sample (samples having any relatives in the dataset using the field 22021: Genetic kinship to other participants) and retricted the study to samples with self-reported British white ancestry (field 21000 with coding 1001). After quality control, we obtained 276,451 individuals and 459,792 SNPs.

## Data availability

The individual-level genotype and phenotype data are available by application from the UKBB http://www.ukbiobank.ac.uk/.

## URLs

SCORE software: https://github.com/sriramlab/SCORE

HDL software is available at https://github.com/zhenin/HDL. The reference panel to run HDL is available at https://github.com/zhenin/HDL/wiki/Reference-panels.

LDSC software is available at https://github.com/bulik/ldsc/.

PLINK1.9 (https://www.cog-genomics.org/plink/2.0) was used to compute summary statistics GCTA-REML and GCTA-HE are available at https://cnsgenomics.com/software/gcta/.

## Acknowledgments

This research was conducted using the UK Biobank Resource under applications 33127 and 33297. We thank the participants of UK Biobank for making this work possible. This work was funded by NIH grants R35GM125055 (S.S.), P01HL28481 (P.P.), and U01DK105561 (P.P.), an Alfred P. Sloan Research Fellowship (S.S.), and NSF grants III-1705121, (Y.W. and S.S).

