## Supplement for "Fast estimation of genetic correlation for Biobank-scale data"

#### SCORE-OVERLAP: Estimating genetic correlation when the traits are measured on the same set of individuals

We describe our model in the setting where the two traits are measured on the same set of individuals and then generalize to the setting where the traits are measured on a partially overlapping set of individuals.

Assuming the two traits overlap across all samples, we let $\boldsymbol{X}$ denote the genotype matrix for the two traits. Let concatenated phenotype vector, $\boldsymbol{y} \equiv [\boldsymbol{y}_1^T, \boldsymbol{y}_2^T]^T$, concatenated environmental effect vector $\boldsymbol{\epsilon} \equiv [\boldsymbol{\epsilon_1}^T, \boldsymbol{\epsilon_2}^T]^T$, and concatenated effect size vector$\boldsymbol{\beta} \equiv [\boldsymbol{\beta_1}^T, \boldsymbol{\beta_2}^T]^T$. We assume the generative model is:

$$\boldsymbol{y} = \begin{bmatrix} \boldsymbol{X} & 0 \\ 0 & \boldsymbol{X} \end{bmatrix} \boldsymbol{\beta} + \boldsymbol{\epsilon} \tag{1}$$

Further, we assume $\mathbb{E}[\boldsymbol{\beta}_1] = 0, \mathbb{E}[\boldsymbol{\beta_2}] = 0$, and have the covariance matrix:

$$\begin{aligned} cov(\boldsymbol{\beta_1}, \boldsymbol{\beta_1}) &= \frac{1}{M}\sigma_{g1}^2 \boldsymbol{I}_M \\ cov(\boldsymbol{\beta_2}, \boldsymbol{\beta_2}) &= \frac{1}{M}\sigma_{g2}^2 \boldsymbol{I}_M \\ cov(\boldsymbol{\beta_1}, \boldsymbol{\beta_2}) &= \frac{1}{M}\gamma_g \boldsymbol{I}_M \end{aligned} \tag{2}$$

We assume $\mathbb{E}[\boldsymbol{y}] = 0$. Thus the population covariance of the concatenated phenotype vector $\boldsymbol{y}$

is given by:

$$cov(\boldsymbol{y}) = \mathbb{E}[\boldsymbol{y}\boldsymbol{y}^T] - \mathbb{E}[\boldsymbol{y}]\mathbb{E}[\boldsymbol{y}]^T = \begin{bmatrix} \sigma_{g1}^2 \boldsymbol{K} & \gamma_g \boldsymbol{K} \\ \gamma_g \boldsymbol{K}^T & \sigma_{g2}^2 \boldsymbol{K} \end{bmatrix} + \begin{bmatrix} \sigma_{e1}^2 I_N & \gamma_e \boldsymbol{I}_N \\ \gamma_e I_N & \sigma_{e2}^2 \boldsymbol{I}_N \end{bmatrix} \tag{3}$$

Here $\boldsymbol{K} = \frac{\boldsymbol{X}\boldsymbol{X}^T}{M}$ is the genetic relatedness matrix (GRM). $\sigma_{gt}^2, \sigma_{et}^2$ denote the genetic and environmental variance components associated with trait $t$. Our approach to estimate both the variance components and the genetic correlation relies on a Method-of-Moments (MoM) estimator obtained by equating the population covariance to the empirical covariance. The empirical covariance of the concatenated phenotype vector $\boldsymbol{y}$ is estimated by the sample covariance: $\boldsymbol{y}\boldsymbol{y}^T$. The MoM estimator is obtained by solving the following ordinary least squares problem:

$$(\widehat{\gamma_g}, \widehat{\gamma_e}, \widehat{\sigma_{g1}^2}, \widehat{\sigma_{g2}^2}, \widehat{\sigma_{e1}^2}, \widehat{\sigma_{e2}^2}) = argmin_{\gamma_g, \gamma_e, \sigma_{g1}^2, \sigma_{g2}^2, \sigma_{e1}^2, \sigma_{e2}^2} ||\boldsymbol{y}\boldsymbol{y}^T - ( \begin{bmatrix} \sigma_{g1}^2 \boldsymbol{K} & \gamma_g \boldsymbol{K} \\ \gamma_g \boldsymbol{K}^T & \sigma_{g2}^2 \boldsymbol{K} \end{bmatrix} + \begin{bmatrix} \sigma_{e1}^2 I_N & \gamma_e I_N \\ \gamma_e I_N & \sigma_{e2}^2 I_N \end{bmatrix} )||_F^2 \tag{4}$$

Setting the gradient of the objective function to zero gives us the normal equations. We observe that solving for the genetic and environmental covariance parameters $(\gamma_g, \gamma_e)$ is decoupled from solving for the variance component parameters: $\sigma_{g1}^2, \sigma_{e1}^2, \sigma_{g2}^2, \sigma_{e2}^2$. Thus, MoM estimates of the covariance parameters can be obtained by solving the set of normal equations:

$$\begin{bmatrix} tr(\boldsymbol{K}^2) & tr(\boldsymbol{K}) \\ tr(\boldsymbol{K}) & N \end{bmatrix} \begin{bmatrix} \widehat{\gamma_g} \\ \widehat{\gamma_e} \end{bmatrix} = \begin{bmatrix} \boldsymbol{y}_2^T \boldsymbol{K} \boldsymbol{y}_1 \\ \boldsymbol{y}_2^T \boldsymbol{y}_1 \end{bmatrix} \tag{5}$$

The GRM $\boldsymbol{K}$ can be computed in time $\mathcal{O}(MN^2)$ and $\mathcal{O}(N^2)$ memory. Given the GRM, computing each of the coefficients for the normal equations requires $\mathcal{O}(N^2)$ time.

Given each of the coefficients, we can solve analytically for $\widehat{\gamma_g}$, and $\widehat{\gamma_e}$:

$$\widehat{\gamma_g} = \frac{\boldsymbol{y}_1^T \boldsymbol{K} \boldsymbol{y}_2 - \boldsymbol{y}_1^T \boldsymbol{y}_2}{tr[\boldsymbol{K}^2] - N}$$

Here we have used the property that $tr(\boldsymbol{K}) = N$ due to the use of a standardized genotype matrix.

Similarly, we solve the following linear systems for the estimators of genetic variances:

$$\begin{bmatrix} tr(\boldsymbol{K}^2) & tr(\boldsymbol{K}) \\ tr(\boldsymbol{K}) & N \end{bmatrix} \begin{bmatrix} \widehat{\sigma^2_{g1}} & \widehat{\sigma^2_{g2}} \\ \widehat{\sigma^2_{e1}} & \widehat{\sigma^2_{e2}} \end{bmatrix} = \begin{bmatrix} \boldsymbol{y}_1^T \boldsymbol{K} \boldsymbol{y}_1 & \boldsymbol{y}_2^T \boldsymbol{K} \boldsymbol{y}_2 \\ \boldsymbol{y}_1^T \boldsymbol{y}_1 & \boldsymbol{y}_2^T \boldsymbol{y}_2 \end{bmatrix} \tag{6}$$

and thus the estimators for $\sigma^2_{g1}$ and $\sigma^2_{g2}$ are give by $\widehat{\sigma^2_{g1}} = \frac{\boldsymbol{y}_1^T \boldsymbol{K} \boldsymbol{y}_1 - \boldsymbol{y}_1^T \boldsymbol{y}_1}{tr[\boldsymbol{K}^2] - N}$ and $\widehat{\sigma^2_{g2}} = \frac{\boldsymbol{y}_2^T \boldsymbol{K} \boldsymbol{y}_2 - \boldsymbol{y}_2^T \boldsymbol{y}_2}{tr[\boldsymbol{K}^2] - N}$

Finally, we use estimates of the genetic variance parameters to obtain a plug-in estimate of the genetic correlation: $\widehat{\rho_g} = \frac{\widehat{\gamma_g}}{\sqrt{\widehat{\sigma^2_{g1}}}\sqrt{\widehat{\sigma^2_{g2}}}}$.

Substituting the expressions for the genetic covariance and variances and the GRM gives us the following estimator of genetic correlation:

$$\widehat{\rho_g} = \frac{\boldsymbol{y}_1^T \boldsymbol{K} \boldsymbol{y}_2 - \boldsymbol{y}_1^T \boldsymbol{y}_2}{\sqrt{\boldsymbol{y}_1^T \boldsymbol{K} \boldsymbol{y}_1 - \boldsymbol{y}_1^T \boldsymbol{y}_1}\sqrt{\boldsymbol{y}_2^T \boldsymbol{K} \boldsymbol{y}_2 - \boldsymbol{y}_2^T \boldsymbol{y}_2}} \tag{7}$$

Computing $\widehat{\rho_g}$ requires computing $\boldsymbol{X}^T \boldsymbol{y}_1$ and $\boldsymbol{X}^T \boldsymbol{y}_2$, and does not require variance components. Using the fact that the genotype matrix only contains entries in $\{0, 1, 2, \}$, we can compute these quantities in time $\mathcal{O}(\frac{NM}{max(\log_3 N, \log_3 M)})$ [1].

### Figures

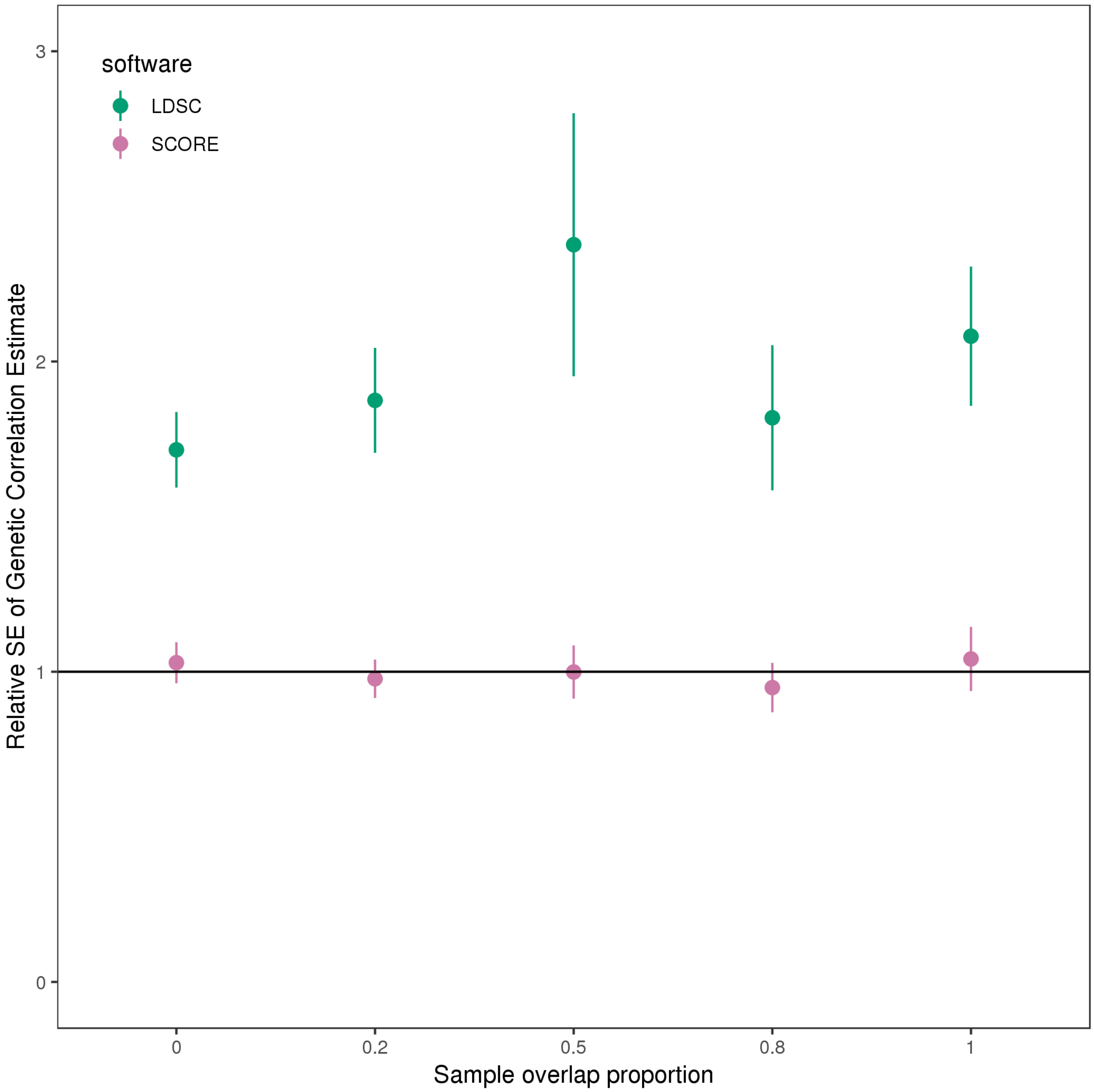


Figure S1: **Comparison of the estimates of genetic correlation from SCORE with GCTA-GREML and LDSC with different proportion of sample overlap ($M = 305,630$ SNPs).** We vary the proportion of sample overlap across the values: $\{0, 0.2, 0.5, 0.8, 0.1\}$. For sample overlap proportion of 0, we have a total of $10,000$ samples where each sample only has observation on one of the traits. For overlap proportion of 1, we have a total $5,000$ samples with each sample having observations on both traits. We report the SE of SCORE and LDSC relative to GCTA-GREML. We ran LDSC with in-sample LD. We estimate the standard error of the relative SE using jackknife.

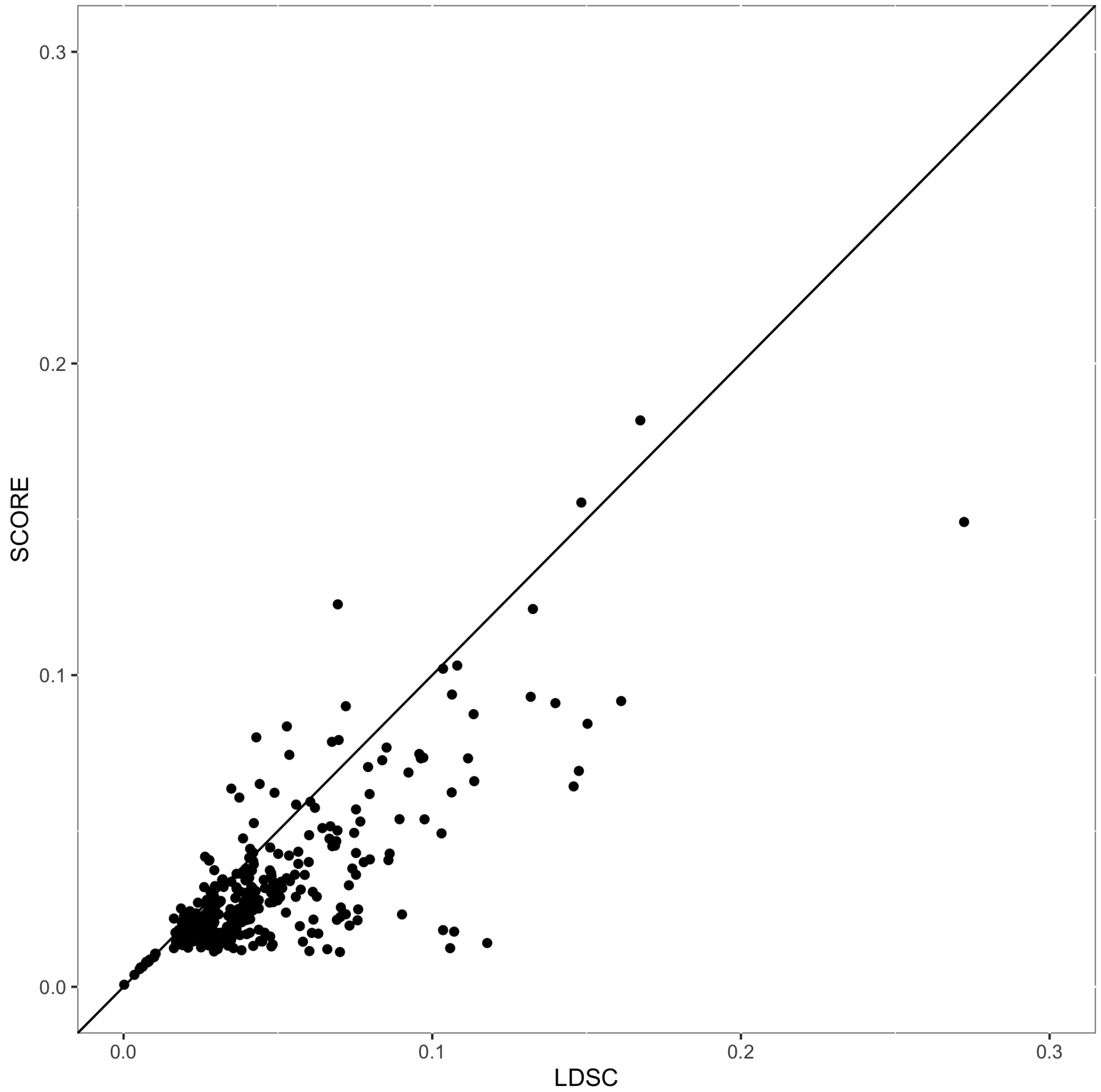


Figure S2: Standard error estimates of genetic correlation between 28 UK biobank phenotypes with LDSC and SCORE corresponding to Figure 4. There are in total 378 tests.

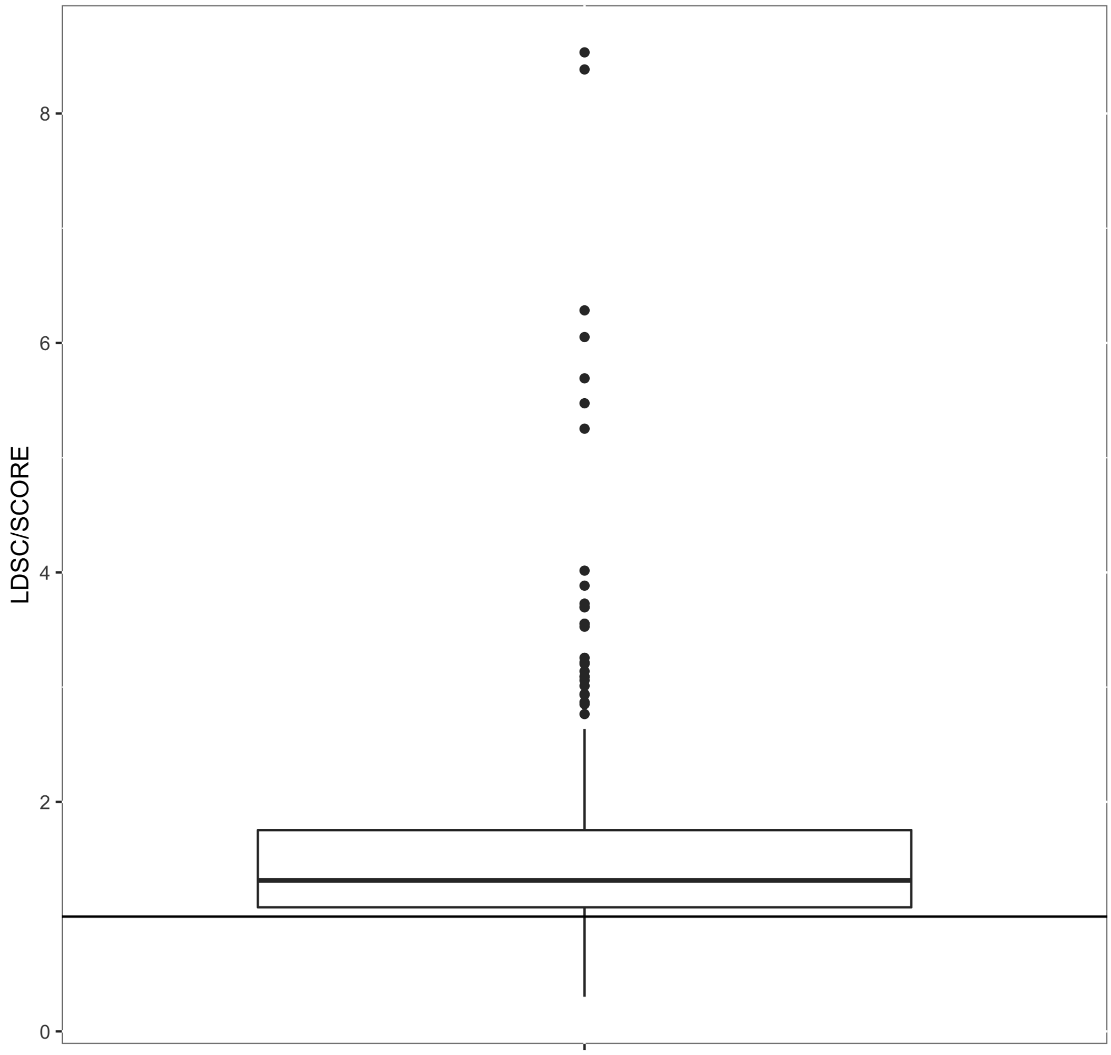


Figure S3: Ratio of standard error estimates of genetic correlation between 28 UK biobank phenotypes with LDSC and SCORE corresponding to Figure 4. There are in total 378 tests.

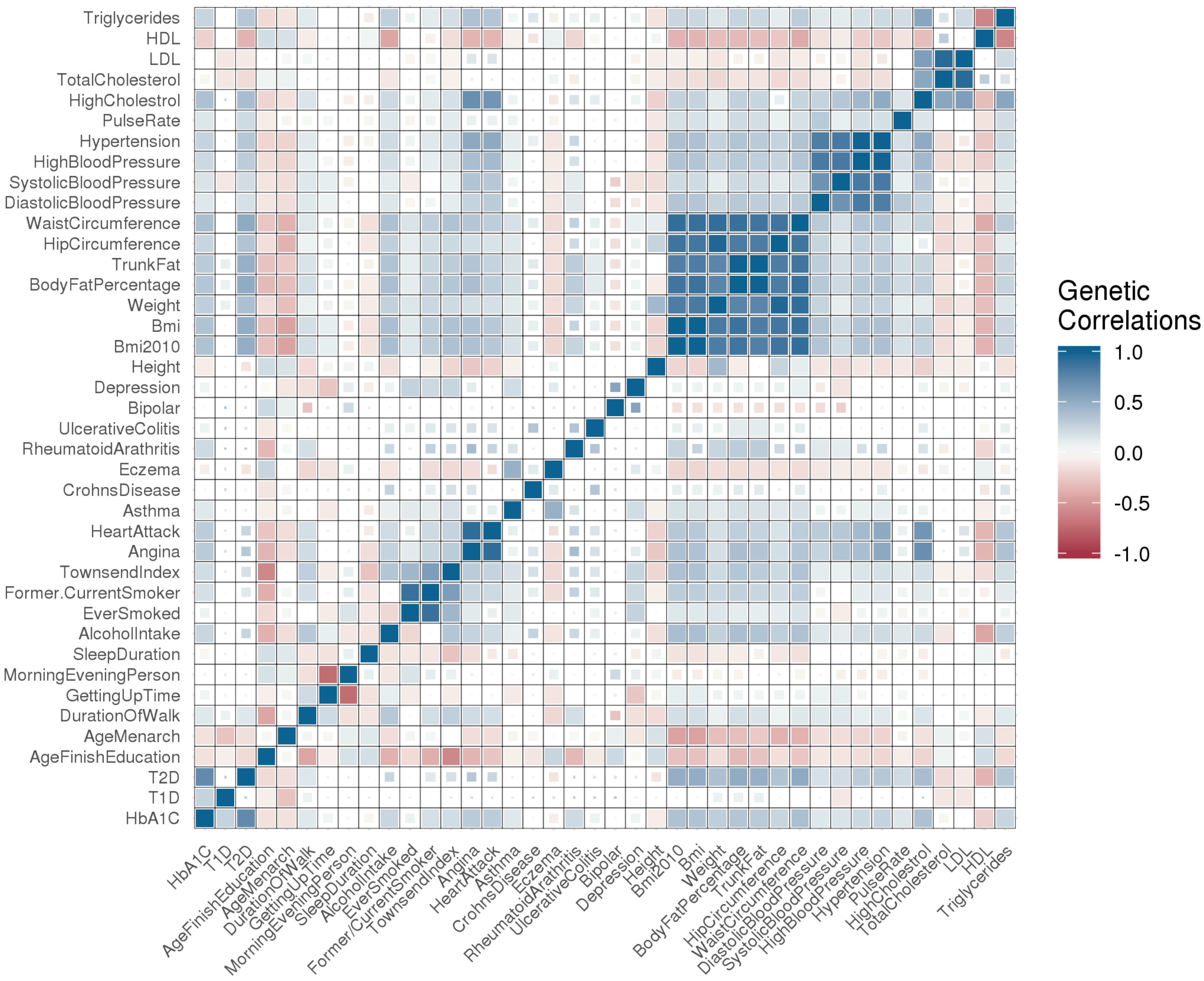


Figure S4: **Genetic correlation estimates in the UK Biobank**: We plot the genetic correlation estimates from SCORE across pairs of 40 phenotypes. Large squares correspond to pairs that are estimated to have a statistically significant genetic correlation after Bonferroni correction at a 5% significance level while small squares indicate pairs that are signficant at a 5% significance level but do not pass multiple-testing correction.

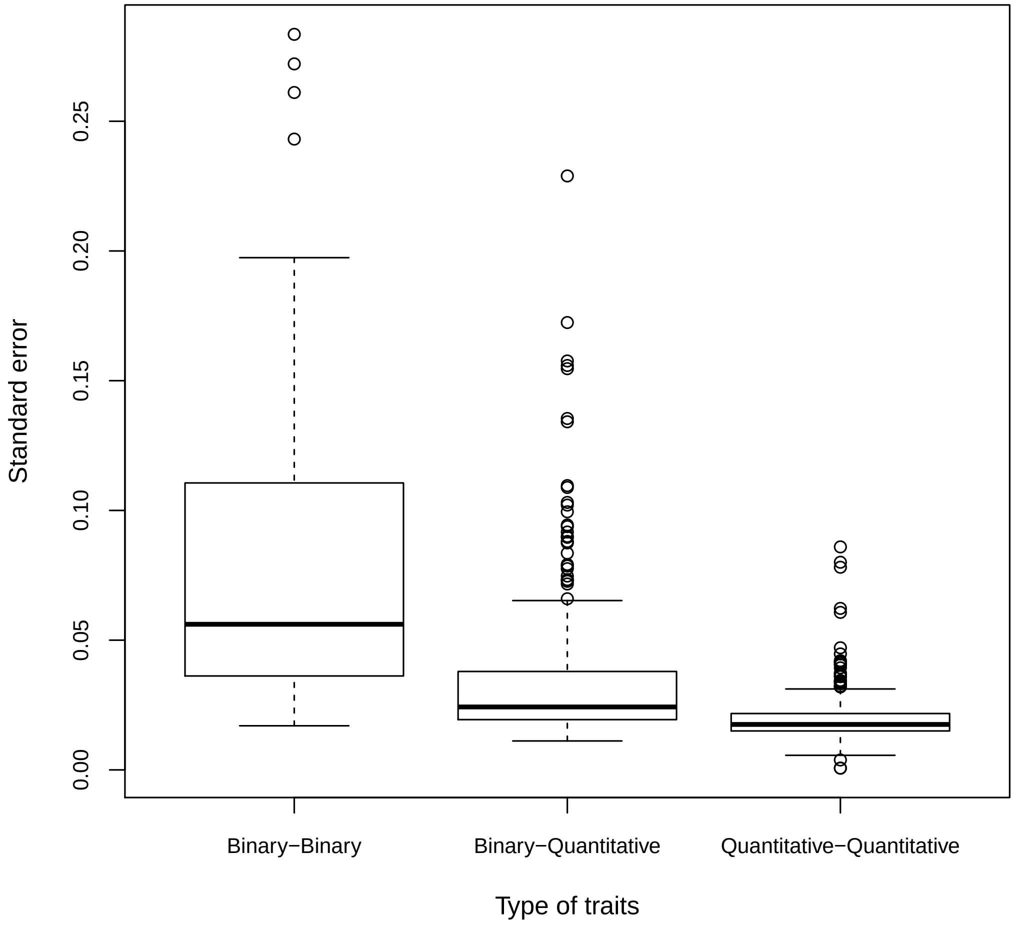


Figure S5: **Standard error of genetic correlation estimates from SCORE by phenotype type**. The three types are: Binary-Binary, Binary-Quantitative, and Quantitative-Quantiative.

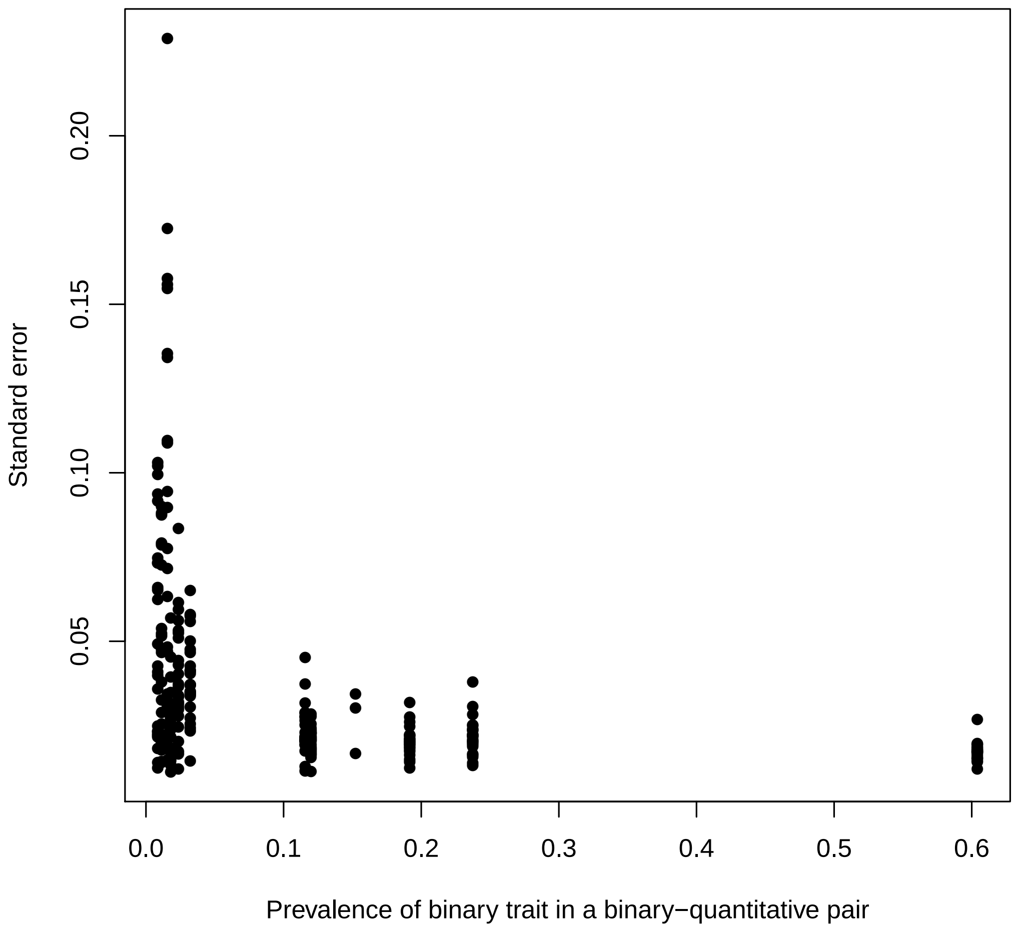


Figure S6: **Standard error of genetic correlation estimates from SCORE by the prevalence of binary phenotype when applied to a Binary-Quantitative pair**.

### Tables

Table S1: **Estimates of bias, mean square error and standard error of SCORE for varying number of random vectors $B = 10$, $B = 100$ and SCORE-OVERLAP**. We simulate a pair of traits with $5,000$ random samples and $305,630$ SNPs from UK Biobank and complete sample overlap. We simulate under both infinitesimal and non-infinitesimal architectures. For non-infinitesimal architectures, the probability of SNPs being causal (having a non-zero effect) for exactly one of the two traits is 0.2 while the probability of a SNP being causal for both traits is 0.1.

| Architecture | Method SCORE | Genetic correlation $\rho_g$ | Bias | MSE | SE | Bias | MSE | SE | Bias | MSE | SE | Bias | MSE | se |
|---|---|---|---|---|---|---|---|---|---|---|---|---|---|---|
| | | | $h_{g1}^2 = 0.1, h_{g2}^2 = 0.2$ | | | $h_{g1}^2 = 0.2, h_{g2}^2 = 0.6$ | | | $h_{g1}^2 = 0.5, h_{g2}^2 = 0.5$ | | | $h_{g1}^2 = 0.6, h_{g2}^2 = 0.8$ | | |
| Infinitesimal | OVERLAP | 0 | -0.0826 | 0.1019 | 0.3083 | -0.0302 | 0.0428 | 0.2048 | 0.0099 | 0.0145 | 0.1201 | 0.0083 | 0.0084 | 0.0912 |
| | B=10 | 0 | -0.0826 | 0.1018 | 0.3082 | -0.0301 | 0.0428 | 0.2047 | 0.0099 | 0.0145 | 0.1201 | 0.0083 | 0.0084 | 0.0912 |
| | B=100 | 0 | -0.0826 | 0.1018 | 0.3082 | -0.0301 | 0.0428 | 0.2047 | 0.0099 | 0.0145 | 0.1201 | 0.0083 | 0.0084 | 0.0912 |
| | OVERLAP | 0.2 | -0.05 | 0.132 | 0.3598 | -0.01 | 0.033 | 0.1814 | 0.0147 | 0.0159 | 0.1253 | -0.0173 | 0.0089 | 0.0928 |
| | B=10 | 0.2 | -0.05 | 0.1319 | 0.3597 | -0.01 | 0.033 | 0.1813 | 0.0147 | 0.0159 | 0.1253 | -0.0173 | 0.0089 | 0.0928 |
| | B=100 | 0.2 | -0.05 | 0.1319 | 0.3597 | -0.01 | 0.033 | 0.1813 | 0.0147 | 0.0159 | 0.1253 | -0.0173 | 0.0089 | 0.0928 |
| | OVERLAP | 0.5 | -0.1567 | 0.0841 | 0.244 | -0.016 | 0.0266 | 0.1622 | 0.0115 | 0.0137 | 0.1164 | 0.0143 | 0.0047 | 0.0669 |
| | B=10 | 0.5 | -0.1568 | 0.0841 | 0.244 | -0.0161 | 0.0266 | 0.1621 | 0.0114 | 0.0137 | 0.1164 | 0.0142 | 0.0047 | 0.0669 |
| | B=100 | 0.5 | -0.1568 | 0.0841 | 0.244 | -0.0161 | 0.0266 | 0.1621 | 0.0116 | 0.0138 | 0.1169 | 0.0142 | 0.0047 | 0.0669 |
| | OVERLAP | 0.8 | -0.3187 | 0.156 | 0.2334 | -0.1106 | 0.0309 | 0.1365 | -0.0206 | 0.0067 | 0.0789 | -0.0016 | 0.0034 | 0.0583 |
| | B=10 | 0.8 | -0.3187 | 0.156 | 0.2333 | -0.1107 | 0.0309 | 0.1365 | -0.0207 | 0.0067 | 0.0789 | -0.0017 | 0.0034 | 0.0583 |
| | B=100 | 0.8 | -0.3187 | 0.156 | 0.2333 | -0.1107 | 0.0309 | 0.1365 | -0.0207 | 0.0067 | 0.0789 | -0.0017 | 0.0034 | 0.0583 |
| Non-Infinitesimal | OVERLAP | 0 | 0.0506 | 0.1227 | 0.3466 | 0.0023 | 0.0349 | 0.1869 | 0.0022 | 0.0198 | 0.1408 | 0.0052 | 0.0091 | 0.0952 |
| | B=10 | 0 | 0.0506 | 0.1226 | 0.3465 | 0.0023 | 0.0349 | 0.1868 | 0.0022 | 0.0198 | 0.1408 | 0.0052 | 0.0091 | 0.0952 |
| | B=100 | 0 | 0.0506 | 0.1226 | 0.3465 | 0.0023 | 0.0349 | 0.1868 | 0.0022 | 0.0198 | 0.1408 | 0.0052 | 0.0091 | 0.0952 |
| | OVERLAP | 0.2 | 0.0259 | 0.0965 | 0.3096 | 0.0116 | 0.0382 | 0.1952 | -6e-04 | 0.0161 | 0.1267 | -0.01 | 0.0065 | 0.0797 |
| | B=10 | 0.2 | 0.0259 | 0.0965 | 0.3096 | 0.0115 | 0.0382 | 0.1952 | -7e-04 | 0.0161 | 0.1267 | -0.01 | 0.0065 | 0.0797 |
| | B=100 | 0.2 | 0.0259 | 0.0965 | 0.3096 | 0.0115 | 0.0382 | 0.1952 | -7e-04 | 0.0161 | 0.1267 | -0.01 | 0.0065 | 0.0797 |
| | OVERLAP | 0.5 | 0.0317 | 0.0974 | 0.3104 | 0.0039 | 0.0287 | 0.1693 | 0.0197 | 0.0118 | 0.1066 | 0.0076 | 0.0083 | 0.0909 |
| | B=10 | 0.5 | 0.0316 | 0.0973 | 0.3104 | 0.0039 | 0.0287 | 0.1693 | 0.0197 | 0.0117 | 0.1066 | 0.0076 | 0.0083 | 0.0909 |
| | B=100 | 0.5 | 0.0316 | 0.0973 | 0.3104 | 0.0039 | 0.0287 | 0.1693 | 0.022 | 0.0114 | 0.1045 | 0.0076 | 0.0083 | 0.0909 |
| | OVERLAP | 0.8 | -0.1181 | 0.1209 | 0.3271 | 0.0116 | 0.031 | 0.1758 | 0.0228 | 0.013 | 0.1119 | 0.031 | 0.0121 | 0.1054 |
| | B=10 | 0.8 | -0.1182 | 0.1209 | 0.327 | 0.0115 | 0.031 | 0.1758 | 0.0228 | 0.013 | 0.1118 | 0.0309 | 0.0121 | 0.1054 |
| | B=100 | 0.8 | -0.1182 | 0.1209 | 0.327 | 0.0115 | 0.031 | 0.1758 | 0.0228 | 0.013 | 0.1118 | 0.0309 | 0.0121 | 0.1054 |

Table S2: **Ratio of SE of summary-statistic methods comparing to SCORE**: The simulations are based on a subset of the UK Biobank with 5000 samples and $305,630$ SNPs. We consider both infinitesimal and non-infinitesimal models. In the non-infinitesimal model, the proportion of causal variants for both traits is 0.3 while the probability that a SNP is causal for exactly one trait is 0.1. The traits are measured on completely overlapping samples. We report the relative statistical efficiency of SCORE versus LDSC and HDL. We also report the relative MSE between SCORE and LDSC, HDL as reference of estimation accuracy.

| Architecture | Method | Genetic correlation $\rho_g$ | 1 | 2 | 3 | 4 |
|---|---|---|---|---|---|---|
| Infinitesimal | LDSC/SCORE | 0 | 1.79 | 1.79 | 2.41 | 2.32 |
| | HDL/SCORE | 0 | 1.35 | 1.2 | 1.42 | 1.35 |
| | LDSC/SCORE | 0.2 | 1.26 | 1.64 | 2.63 | 2.32 |
| | HDL /SCORE | 0.2 | 1.05 | 1.48 | 1.47 | 1.5 |
| | LDSC/SCORE | 0.5 | 1.69 | 2 | 2.19 | 2.6 |
| | HDL/SCORE | 0.5 | 1.65 | 1.14 | 1.18 | 1.39 |
| | LDSC/SCORE | 0.8 | 1.4 | 1.62 | 1.95 | 2.16 |
| | HDL/SCORE | 0.8 | 1.26 | 1.23 | 1.41 | 1.28 |
| Non-Infinitesimal | LDSC/SCORE | 0 | 1.4 | 1.96 | 2.08 | 2.07 |
| | HDL/SCORE | 0 | 1.25 | 1.41 | 1.21 | 1.36 |
| | LDSC/SCORE | 0.2 | 1.74 | 2.09 | 2.15 | 2.28 |
| | HDL/SCORE | 0.2 | 1.44 | 1.54 | 1.38 | 1.4 |
| | LDSC/SCORE | 0.5 | 1.49 | 1.96 | 2.59 | 2.37 |
| | HDL/SCORE | 0.5 | 1.41 | 1.32 | 1.54 | 1.42 |
| | LDSC/SCORE | 0.8 | 1.6 | 2.01 | 2.42 | 2.2 |
| | HDL/SCORE | 0.8 | 1.17 | 1.45 | 1.61 | 1.23 |

Table S3: **Estimates of bias, mean square error and standard error of genetic correlation estimation methods in simulations corresponding to Fig 1**. We simulate a pair of traits with $5,000$ random samples and $305,630$ SNPs from the UK Biobank and complete sample overlap. We simulate under the infinitesimal model where all genetic variants have a small effect on both traits and vary trait heritability and genetic correlation.

| Method | Genetic correlation $\rho_g$ | Bias | MSE | SE | Bias | MSE | SE | Bias | MSE | SE | Bias | MSE | SE |
|---|---|---|---|---|---|---|---|---|---|---|---|---|---|
| | | $h^2_{g1} = 0.1, h^2_{g2} = 0.2$ | | | $h^2_{g1} = 0.2, h^2_{g2} = 0.6$ | | | $h^2_{g1} = 0.5, h^2_{g2} = 0.5$ | | | $h^2_{g1} = 0.6, h^2_{g2} = 0.8$ | | |
| GCTA-GREML | 0 | -0.0818 | 0.0998 | 0.305 | -0.029 | 0.0395 | 0.197 | 0.0096 | 0.013 | 0.114 | 0.0014 | 0.0076 | 0.0871 |
| GCTA-HE | 0 | -0.104 | 0.217 | 0.454 | 0.0366 | 0.0878 | 0.294 | 0.0026 | 0.0268 | 0.164 | 0.0225 | 0.0148 | 0.119 |
| HDL | 0 | -0.0265 | 0.172 | 0.414 | -0.0095 | 0.0606 | 0.246 | 0.0185 | 0.0294 | 0.17 | 0.007 | 0.0153 | 0.123 |
| LDSC | 0 | 0.0519 | 0.3055 | 0.5503 | -0.007 | 0.1343 | 0.3665 | -0.0014 | 0.0837 | 0.2894 | 0.0083 | 0.045 | 0.2119 |
| SCORE | 0 | -0.0826 | 0.102 | 0.308 | -0.0302 | 0.0428 | 0.205 | 0.0099 | 0.0145 | 0.12 | 0.0083 | 0.0084 | 0.0912 |
| GCTA-GREML | 0.2 | -0.0571 | 0.123 | 0.346 | -0.0103 | 0.0328 | 0.181 | 0.0074 | 0.0152 | 0.123 | -0.0114 | 0.0077 | 0.0868 |
| GCTA-HE | 0.2 | -0.188 | 0.261 | 0.475 | -0.0234 | 0.0554 | 0.234 | 0.029 | 0.0321 | 0.177 | -0.0263 | 0.0157 | 0.123 |
| HDL | 0.2 | -0.094 | 0.151 | 0.377 | 0.0254 | 0.0729 | 0.269 | 0.0236 | 0.0346 | 0.184 | 0.0027 | 0.0194 | 0.139 |
| LDSC | 0.2 | -0.0946 | 0.2134 | 0.4522 | 0.0134 | 0.0885 | 0.2973 | 0.0177 | 0.1088 | 0.3293 | 0.0051 | 0.0463 | 0.215 |
| SCORE | 0.2 | -0.05 | 0.132 | 0.36 | -0.01 | 0.033 | 0.181 | 0.0147 | 0.0159 | 0.125 | -0.0173 | 0.0089 | 0.0928 |
| GCTA-GREML | 0.5 | -0.158 | 0.0858 | 0.247 | -0.012 | 0.0244 | 0.156 | 0.0159 | 0.0121 | 0.109 | 0.0092 | 0.0045 | 0.0662 |
| GCTA-HE | 0.5 | -0.142 | 0.161 | 0.375 | -0.0394 | 0.0593 | 0.24 | 0.0181 | 0.0224 | 0.149 | 0.0111 | 0.0116 | 0.107 |
| HDL | 0.5 | -0.162 | 0.188 | 0.401 | -0.0195 | 0.0345 | 0.185 | 0.0047 | 0.0189 | 0.138 | 0.0207 | 0.009 | 0.0928 |
| LDSC | 0.5 | -0.3298 | 0.278 | 0.4114 | -0.0955 | 0.1143 | 0.3243 | 0 | 0.0649 | 0.2547 | 0.0308 | 0.0312 | 0.1739 |
| SCORE | 0.5 | -0.1567 | 0.0841 | 0.244 | -0.016 | 0.0266 | 0.1622 | 0.0115 | 0.0137 | 0.1164 | 0.0143 | 0.0047 | 0.0669 |
| GCTA-GREML | 0.8 | -0.317 | 0.156 | 0.235 | -0.106 | 0.0283 | 0.131 | -0.0232 | 0.0073 | 0.082 | -0.0004 | 0.0033 | 0.0571 |
| GCTA-HE | 0.8 | -0.389 | 0.287 | 0.369 | -0.115 | 0.0446 | 0.177 | -0.023 | 0.0139 | 0.116 | 0.0093 | 0.0061 | 0.0778 |
| HDL | 0.8 | -0.391 | 0.239 | 0.294 | -0.154 | 0.0518 | 0.168 | -0.0421 | 0.0141 | 0.111 | -0.0114 | 0.0057 | 0.0748 |
| LDSC | 0.8 | -0.4841 | 0.3415 | 0.3273 | -0.1851 | 0.0833 | 0.2215 | -0.0727 | 0.029 | 0.1541 | -0.019 | 0.0162 | 0.1257 |
| SCORE | 0.8 | -0.319 | 0.156 | 0.233 | -0.111 | 0.0309 | 0.137 | -0.0206 | 0.0067 | 0.0789 | -0.0016 | 0.0034 | 0.0583 |

Table S4: **Estimates of bias, mean square error and standard error of genetic correlation estimation methods in simulations corresponding to Fig 2**. We simulate a pair of traits with 5,000 random samples and 305,630 SNPs from UK Biobank and complete sample overlap. We simulate under a non-infinitesimal model with the proportion of SNPs being causal for exactly one of the two traits is 0.2 and the probability of a SNP being causal on both traits is 0.1. We vary the heritability of each trait and genetic correlation.

| Method | Genetic correlation $\rho_g$ | Bias | MSE | SE | Bias | MSE | SE | Bias | MSE | SE | Bias | MSE | SE |
|---|---|---|---|---|---|---|---|---|---|---|---|---|---|
| | | $h^2_{g1} = 0.1, h^2_{g2} = 0.2$ | | | $h^2_{g1} = 0.2, h^2_{g2} = 0.6$ | | | $h^2_{g1} = 0.5, h^2_{g2} = 0.5$ | | | $h^2_{g1} = 0.6, h^2_{g2} = 0.8$ | | |
| GCTA-GREML | 0 | 0.0414 | 0.12 | 0.344 | 0.003 | 0.0343 | 0.185 | -0.0025 | 0.0157 | 0.125 | -0.0011 | 0.0075 | 0.0866 |
| GCTA-HE | 0 | 0.0964 | 0.243 | 0.484 | 0.0163 | 0.0502 | 0.223 | 0.0103 | 0.0328 | 0.181 | -0.0051 | 0.0159 | 0.126 |
| HDL | 0 | 0.0626 | 0.193 | 0.434 | -0.0225 | 0.0699 | 0.263 | 0.0103 | 0.0292 | 0.171 | 0.0155 | 0.0171 | 0.13 |
| LDSC | 0 | 0.0608 | 0.2375 | 0.4835 | -0.0273 | 0.1344 | 0.3656 | 0.0179 | 0.0862 | 0.2931 | 0.0301 | 0.0399 | 0.1975 |
| SCORE | 0 | 0.0506 | 0.123 | 0.347 | 0.0023 | 0.0349 | 0.187 | 0.0022 | 0.0198 | 0.141 | 0.0052 | 0.0091 | 0.0952 |
| GCTA-GREML | 0.2 | -0.0034 | 0.12 | 0.347 | 0.0143 | 0.0376 | 0.193 | -0.0055 | 0.0159 | 0.126 | -0.007 | 0.0058 | 0.0759 |
| GCTA-HE | 0.2 | 0.0162 | 0.2 | 0.447 | 0.0182 | 0.0706 | 0.265 | -0.0073 | 0.0298 | 0.172 | -0.0247 | 0.0168 | 0.127 |
| HDL | 0.2 | -0.0066 | 0.198 | 0.445 | 0.0155 | 0.0902 | 0.3 | 0.005 | 0.0307 | 0.175 | -0.0121 | 0.0125 | 0.111 |
| LDSC | 0.2 | -0.0846 | 0.2979 | 0.5392 | 0.0209 | 0.1664 | 0.4073 | -0.0089 | 0.0744 | 0.2726 | 0.0052 | 0.033 | 0.1815 |
| SCORE | 0.2 | 0.0259 | 0.0965 | 0.31 | 0.0116 | 0.0382 | 0.195 | -0.0006 | 0.0161 | 0.127 | -0.01 | 0.0065 | 0.0797 |
| GCTA-GREML | 0.5 | 0.0292 | 0.102 | 0.319 | 0.0081 | 0.0289 | 0.17 | 0.0182 | 0.0121 | 0.108 | 0.0068 | 0.0075 | 0.0861 |
| GCTA-HE | 0.5 | -0.0213 | 0.146 | 0.382 | 0.0061 | 0.0576 | 0.24 | 0.0138 | 0.0226 | 0.15 | 0.0155 | 0.0155 | 0.123 |
| HDL | 0.5 | 0.0137 | 0.124 | 0.352 | -0.0255 | 0.0552 | 0.234 | 0.0223 | 0.0304 | 0.173 | 0.0055 | 0.0167 | 0.129 |
| LDSC | 0.5 | -0.1026 | 0.2247 | 0.4628 | -0.0634 | 0.1138 | 0.3313 | -0.0142 | 0.0766 | 0.2764 | -0.0202 | 0.0467 | 0.2151 |
| SCORE | 0.5 | 0.0317 | 0.0974 | 0.31 | 0.0039 | 0.0287 | 0.169 | 0.0197 | 0.0118 | 0.107 | 0.0076 | 0.0083 | 0.0909 |
| GCTA-GREML | 0.8 | -0.112 | 0.12 | 0.328 | 0.0146 | 0.0301 | 0.173 | 0.0164 | 0.0134 | 0.115 | 0.036 | 0.0095 | 0.0904 |
| GCTA-HE | 0.8 | -0.0941 | 0.212 | 0.451 | 0.0307 | 0.0454 | 0.211 | 0.0151 | 0.0277 | 0.166 | 0.0288 | 0.0185 | 0.133 |
| HDL | 0.8 | -0.155 | 0.172 | 0.385 | 0.0355 | 0.0665 | 0.255 | 0.0362 | 0.0336 | 0.18 | 0.0453 | 0.019 | 0.13 |
| LDSC | 0.8 | -0.2322 | 0.3293 | 0.5248 | 0.0022 | 0.125 | 0.3536 | 0.0144 | 0.0733 | 0.2704 | 0.0134 | 0.0542 | 0.2324 |
| SCORE | 0.8 | -0.118 | 0.121 | 0.327 | 0.0116 | 0.031 | 0.176 | 0.0228 | 0.013 | 0.112 | 0.031 | 0.0121 | 0.105 |

Table S5: **SCORE vs. LDSC varying proportion of sample overlap**. The bias, MSE and variance corresponding to Figure S1. Given the true genetic correlation being 0.5 and the heritability of pair of traits being $\{0.2, 0.6\}$, we vary the sample overlap sample across $\{0, 0.2, 0.5, 0.8, 1\}$.

| Overlap Proportion | Software | bias | MSE | SE |
|---|---|---|---|---|
| 0 | LDSC | -0.1672 | 0.1405 | 0.3355 |
| 0.2 | LDSC | 0.0044 | 0.0921 | 0.3035 |
| 0.5 | LDSC | 0.1708 | 0.1022 | 0.2702 |
| 0.8 | LDSC | 0.0808 | 0.0815 | 0.2737 |
| 1 | LDSC | -0.0955 | 0.1143 | 0.3243 |
| 0 | SCORE | -0.0037 | 0.0405 | 0.2013 |
| 0.2 | SCORE | 0.2242 | 0.0753 | 0.1582 |
| 0.5 | SCORE | 0.2843 | 0.0916 | 0.1036 |
| 0.8 | SCORE | 0.2722 | 0.087 | 0.1428 |
| 1 | SCORE | -0.016 | 0.0266 | 0.1622 |
| 0 | GCTA-GREML | -0.0063 | 0.0383 | 0.1956 |
| 0.2 | GCTA-GREML | 0.2635 | 0.0957 | 0.1619 |
| 0.5 | GCTA-GREML | 0.3301 | 0.1219 | 0.1137 |
| 0.8 | GCTA-GREML | 0.2371 | 0.0788 | 0.1505 |
| 1 | GCTA-GREML | -0.012 | 0.0244 | 0.1558 |

Table S6: **Jackknife estimates of standard error are accurate**. The simulations are based on a subset of UK Biobank with $5,000$ samples and $305,630$ SNPs. We consider infinitesimal and non-infinitesimal architectures with varying heritability and genetic correlation. In the non-infinitesimal model, the probability of variants being causal for both traits is 0.1 and the probability of a variant being causal for exactly one of the two traits is 0.2. We consider the case of complete sample overlap. We report the average of estimates of standard error across 100 trials. We use block size of 4000 SNPs for the Jackknife.

| Architecture | Heritability | $\rho_g$ | $\hat{se}$ | se |
|---|---|---|---|---|
| Infinitesimal | 0.1, 0.2 | 0 | 0.45 | 0.42 |
| | 0.2, 0.6 | 0 | 0.2 | 0.2 |
| | 0.5, 0.5 | 0 | 0.12 | 0.12 |
| | 0.6, 0.8 | 0 | 0.09 | 0.09 |
| | 0.1, 0.2 | 0.2 | 0.4 | 0.38 |
| | 0.2, 0.6 | 0.2 | 0.19 | 0.19 |
| | 0.5, 0.5 | 0.2 | 0.12 | 0.12 |
| | 0.6, 0.8 | 0.2 | 0.09 | 0.09 |
| | 0.1, 0.2 | 0.5 | 0.41 | 0.3 |
| | 0.2, 0.6 | 0.5 | 0.18 | 0.17 |
| | 0.5, 0.5 | 0.5 | 0.11 | 0.11 |
| | 0.6, 0.8 | 0.5 | 0.07 | 0.07 |
| | 0.1, 0.2 | 0.8 | 0.41 | 0.34 |
| | 0.2, 0.6 | 0.8 | 0.18 | 0.15 |
| | 0.5, 0.5 | 0.8 | 0.09 | 0.09 |
| | 0.6, 0.8 | 0.8 | 0.05 | 0.05 |
| Non-Infinitesimal | 0.1, 0.2 | 0 | 0.45 | 0.39 |
| | 0.2, 0.6 | 0 | 0.19 | 0.19 |
| | 0.5, 0.5 | 0 | 0.12 | 0.14 |
| | 0.6, 0.8 | 0 | 0.09 | 0.09 |
| | 0.1, 0.2 | 0.2 | 0.43 | 0.39 |
| | 0.2, 0.6 | 0.2 | 0.18 | 0.19 |
| | 0.5, 0.5 | 0.2 | 0.12 | 0.12 |
| | 0.6, 0.8 | 0.2 | 0.09 | 0.08 |
| | 0.1, 0.2 | 0.5 | 0.43 | 0.36 |
| | 0.2, 0.6 | 0.5 | 0.18 | 0.18 |
| | 0.5, 0.5 | 0.5 | 0.12 | 0.11 |
| | 0.6, 0.8 | 0.5 | 0.09 | 0.09 |
| | 0.1, 0.2 | 0.8 | 0.44 | 0.36 |
| | 0.2, 0.6 | 0.8 | 0.20 | 0.18 |
| | 0.5, 0.5 | 0.8 | 0.12 | 0.11 |
| | 0.6, 0.8 | 0.8 | 0.1 | 0.11 |

Table S7: Estimates of genetic correlation as a function of the prevalence of binary traits. We simulate a pair of traits where one of the traits is binary. We also simulate assuming different values of the environmental correlation (no environmenta correlation, the same sign and opposite sign as the genetic correlation). We observe that SCORE has relatively stable SE when the prevalence is greater than 0.5%

| $\sigma^2_{g1}$ | $\sigma^2_{g2}$ | $\rho_g$ | $\rho_e$ | Prevalence | $\widehat{\rho_g}$ | SE |
|---|---|---|---|---|---|---|
| 0.272 | 0.12 | -0.23 | 0 | Continuous trait | -0.238 | 0.093 |
| 0.272 | 0.12 | -0.23 | 0 | 50% | -0.238 | 0.097 |
| 0.272 | 0.12 | -0.23 | 0 | 25% | -0.239 | 0.101 |
| 0.272 | 0.12 | -0.23 | 0 | 10% | -0.238 | 0.103 |
| 0.272 | 0.12 | -0.23 | 0 | 1% | -0.234 | 0.124 |
| 0.272 | 0.12 | -0.23 | 0 | 0.05% | -0.248 | 0.134 |
| 0.272 | 0.12 | -0.23 | 0 | 0.01% | -0.215 | 0.205 |
| 0.272 | 0.12 | -0.23 | -0.04 | Continuous trait | -0.211 | 0.107 |
| 0.272 | 0.12 | -0.23 | -0.04 | 0.01% | -0.332 | 0.386 |
| 0.272 | 0.12 | -0.23 | 0.04 | Continuous trait | -0.241 | 0.079 |
| 0.272 | 0.12 | -0.23 | 0.04 | 0.01% | -0.235 | 0.352 |

Table S8: **Group of 40 traits in UK Biobank**

| | |
|---|---|
| Glucose metabolism and diabetes traits | HbA1C |
| Glucose metabolism and diabetes traits | T1D |
| Glucose metabolism and diabetes traits | T2D |
| Socioeconomic and general medical information traits | Age Finish Education |
| Socioeconomic and general medical information traits | Age Menarch |
| Socioeconomic and general medical information traits | Duration Of Walk |
| Socioeconomic and general medical information traits | Getting Up Time |
| Socioeconomic and general medical information traits | Morning Evening Person |
| Socioeconomic and general medical information traits | Sleep Duration |
| Environmental factor traits | Alcohol Intake |
| Environmental factor traits | Ever Smoked |
| Environmental factor traits | Former/Current Smoker |
| Environmental factor traits | Townsend Index |
| Coronary artery disease related traits | Angina |
| Coronary artery disease related traits | Heart Attack |
| Autoimmune disorders | Asthma |
| Autoimmune disorders | Crohn's Disease |
| Autoimmune disorders | Eczema |
| Autoimmune disorders | Rheumatoid Arthritis |
| Autoimmune disorders | Ulcerative Colitis |
| Psychiatric disorders | Bipolar |
| Psychiatric disorders | Depression |
| Anthropometric traits | Bmi2010 |
| Anthropometric traits | Bmi |
| Anthropometric traits | Body Fat Percentage |
| Anthropometric traits | Height |
| Anthropometric traits | Hip Circumference |
| Anthropometric traits | Trunk Fat |
| Anthropometric traits | Weight |
| Anthropometric traits | Waist Circumference |
| Blood pressure and circulatory traits | Diastolic Blood Pressure |
| Blood pressure and circulatory traits | High Blood Pressure |
| Blood pressure and circulatory traits | Hypertension |
| Blood pressure and circulatory traits | Pulse Rate |
| Blood pressure and circulatory traits | Systolic Blood Pressure |
| Lipid metabolism traits | HDL |
| Lipid metabolism traits | High Cholesterol |
| Lipid metabolism traits | LDL |
| Lipid metabolism traits | Total Cholesterol |
| Lipid metabolism traits | Triglycerides |
